# Systematic Classification Differences Across Eye-Movement Detection Algorithms

**DOI:** 10.1101/2025.09.16.676657

**Authors:** Jonathan Nir, Leon Y. Deouell

**Author notes:** **Author Notes:** JN built the pEYES package, performed analyses and wrote the paper. JN and LYD reviewed the manuscript and prepared it for publication. Both authors approved the final version of the manuscript for submission. All data and code are made available in https://github.com/huji-hcnl/pEYES. Correspondence should be addressed to Jonathan Nir, Edmond and Lily Safra Center for Brain Sciences, The Hebrew University of Jerusalem, Jerusalem 9190401, Israel,.

## Abstract

Eye movement (EM) detection is a critical step in most eye-tracking (ET) research, typically relying on *detectors* – specialized algorithms designed to segment raw ET data into discrete oculomotor events. However, variability in detection algorithms and the lack of standardized evaluation frameworks hinder transparency and reproducibility across studies. In this work, we introduce *pEYES*, an open-source toolkit designed to streamline EM detection and enable robust, quantitative comparisons between detectors. The toolkit provides implementations for several widely used threshold-based detectors, along with multiple standardized evaluation procedures for assessing detection performance.

Using *pEYES*, we evaluated seven detection algorithms on two publicly-available human-annotated datasets containing recordings of subjects freely viewing color images. Performance was assessed using metrics such as Cohen’s Kappa, Relative Timing Offset and Deviation, and a sensitivity index (𝑑′) for fixation and saccade onsets and offsets. *Engbert’s* adaptive velocity-threshold algorithm consistently matched or outperformed the other detectors, occasionally achieving human-level precision. In contrast, several other detectors exhibited substantial variability in performance between datasets. We also found systematic differences in detection scores between fixation and saccade boundaries, with fixation offsets and saccade onsets detected more reliably than their counterparts. These findings highlight the importance of task- and dataset-specific detector selection in EM analysis.

The *pEYES* toolkit is freely available, and its codebase – along with the analyses presented in this report – is accessible at https://github.com/huji-hcnl/pEYES. We invite the research community to use, extend, and contribute to its ongoing development. Through open collaboration, we aim to advance the rigor and reproducibility of EM detection practices.

## Introduction

In recent years, eye-tracking (ET) technology has become increasingly prevalent, revolutionizing research and applications across multiple disciplines (Alhilo & Al-Sakaa, 2023). In particular, ET plays a crucial role in cognitive research, providing invaluable insights into areas like perception and psychophysics (Lima & Ventura, 2023), memory (Schmidig et al., 2025), and applications in clinical neuroscience (Wolf et al., 2023).

Modern ET devices use fast video recording to detect the position of the pupil and corneal reflection, and, following calibration with targets of known positions, compute the location of gaze. The resulting raw data consists of timeseries of estimated gaze coordinates. To make this data more interpretable, it is typically segmented into discrete oculomotor events through a process known as **event detection^1^** – assigning each sample a specific label representing a *type* of eye movement (EM). This classification is essential for linking oculomotor behavior to perceptual, cognitive, or other neural processes, enabling researchers to quantify phenomena such as attention allocation, reading dynamics, or visual search patterns (Kasneci et al., 2024).

In defining events, we use a functional operationalization of the different EM types (Hessels et al., 2018), characterized by their behavioral purpose and features in the ET signal: ***fixations*** are periods of time when gaze remains relatively stable on an area of interest, enabling visual intake on a part of the stimulus (Dar et al., 2021; Kasneci et al., 2014). On the other hand, ***saccades*** are short, high-velocity ballistic movements that shift the position of gaze to a target region (Dar et al., 2021). Unlike saccades, ***smooth pursuits*** (SPs) involve slow and continuous movements of the gaze, which enables visual intake when tracking a moving target (Dar et al., 2021; Kasneci et al., 2024). Additionally, ***post-saccadic oscillations*** (PSOs) are short periods of oculomotor instability that immediately follow some, though not all, saccades (Nyström & Holmqvist, 2010). Lastly, ***blinks*** are defined as short periods of time that include rapid closing and re-opening of the eyelids (Nyström et al., 2024), resulting in temporary loss of ET signal.

The process of EM detection – i.e., assigning labels to data samples – can theoretically be done manually; however, as noted by Komogortsev et al. (2010), this process is “extremely tedious and time-consuming”. Additionally, subjective definitions can result in inconsistent movement classifications, impeding replicability (Hessels et al., 2018). Therefore, researchers typically employ automated algorithms, or *“detectors”*, to assign EM labels systematically, streamlining the process and ensuring consistency.

Although these detectors aim to serve the same purpose, they differ significantly in fundamental characteristics. First, commonly used detectors provided by eye-tracker manufacturers are often proprietary, limiting transparency and the ability to independently review their methods. By contrast, open-source algorithms are transparent, but often have varied implementations across research facilities, undermining reproducibility. Second, as modern computers advance, newer detectors become more complex, requiring more runtime and computational resources. Third, detectors vary in the types of movements they can detect, from basic algorithms that differentiate only between fixations and saccades, to more complex detectors that identify a range of movement types, including PSOs, SPs, and other nuanced EMs. Lastly, detectors also employ various identification techniques, ranging from single-threshold-based algorithms (Komogortsev & Karpov, 2013; Salvucci & Goldberg, 2000), to adaptive-threshold approaches (Dar et al., 2021; Engbert & Kliegl, 2003; Engbert & Mergenthaler, 2006; Nyström & Holmqvist, 2010), probabilistic methods (Kasneci et al., 2014; Santini et al., 2016), and modern machine-learning and deep-learning based models (Fuhl et al., 2021; Startsev et al., 2019; Zemblys et al., 2018).

These variations across algorithms may yield diverse event classifications for the same raw data, which, in turn, raises the important question of how to select the optimal detector. Naturally, technical aspects like runtime, computational efficiency, and application-specific factors – such as the types of eye movements the algorithm can and cannot detect – should be the primary considerations in making this decision. However, many detectors share seemingly similar technical features, or there may be different versions of the same detector with varying hyperparameter configurations, making performance evaluation essential.

Here, we provide a comparative evaluation of threshold-based EM detection algorithms. As demonstrated by Andersson et al. (2017), there is no “one algorithm to rule them all”, as detector performance varies based on the presented stimulus, the behavioral task (e.g., free-viewing vs. object tracking), and the types of EMs the detector can detect. Therefore, we focused our evaluation on detecting fixations and saccades during free-viewing of image stimuli. However, to facilitate reproducibility and extend application to other tasks and stimuli, we introduce ***pEYES*** – an open-source toolkit for simplifying EM detection and quantitatively comparing detection algorithms. Using *pEYES*, we applied several commonly used detectors to a subset of image trials from Andersson et al.’s (2017) *lund2013* dataset. We compared each detector’s output with the human-annotated labels provided in the dataset, and quantified their performance using multiple evaluation procedures and metrics. Our analyses revealed significant performance differences across detectors, with some algorithms consistently aligning more closely with human annotations than others. These findings underscore the need for standardized, task-specific evaluations when selecting a detection algorithm for EM analysis.

## Methods

### The pEYES Package

To evaluate EM detection algorithms, we developed *pEYES*, a toolkit written in the Python programming language (Van Rossum & Drake, 1995), intended for scientific use by ET researchers. It serves two primary goals: (1) to standardize and simplify EM detection; and (2) to provide a framework for evaluating detection performance across multiple algorithms.

To support the first goal, *pEYES* incorporates principles of object-oriented-programming (OOP), to define key concepts of the EM detection process: the `Detector` and the `Event`. A `Detector` represents an instance of EM detection algorithm, configured with a set of user-defined hyperparameters. The package includes a set of pre-implemented threshold-based detectors (listed in **Table 1**), as well as an API for integrating new, custom-implemented detectors.. Similarly, an `Event` object represents a single EM event – a sequence of consecutive samples sharing the same EM type – and precompute its features based on the underlying sample data. *pEYES* also provides commonly-used visualizations to support ET data analysis, whether for a single recording session or for exploring features of specific EM types (see examples in **Appendix A**).

**Table 1:** Detectors implemented in pEYES.

| Algorithm | Threshold Type | Ref. | EM Type |  |  |  |  |
| --- | --- | --- | --- | --- | --- | --- | --- |
|  |  |  | FIX | SAC | PSO | SP | BLI |
| I-VT | Global | Salvucci & Goldberg, 2000 | ✓ | ✓ |  |  | ✓ |
| I-DT | Global | Salvucci & Goldberg, 2000 | ✓ | ✓ |  |  | ✓ |
| I-VVT | Global | Komogortsev & Karpov, 2013 | ✓ | ✓ |  | ✓ | ✓ |
| I-DVT | Global | Komogortsev & Karpov, 2013 | ✓ | ✓ |  | ✓ | ✓ |
| Engbert | Adaptive | Engbert & Kliegl, 2003;<br>Engbert & Mergenthaler, 2006 | ✓ | ✓ |  |  | ✓ |
| NH | Adaptive | Nyström & Holmqvist, 2010 | ✓ | ✓ | ✓ |  | ✓ |
| REMoDNaV | Adaptive | Dar et al., 2021 | ✓ | ✓ | ✓ | ✓ | ✓ |

*Note:* List of detectors available in *pEYES* and the types of EMs they are designed to detect: fixations (FIX), saccades (SAC), post-saccadic oscillations (PSO), smooth pursuits (SP), and blinks (BLI).

To promote the second goal, *pEYES* includes a framework for evaluating detection performance across multiple algorithms. This evaluation can be conducted qualitatively, using dedicated visualizations to compare EM features across detectors (for examples, see **Figure 1**); or quantitatively, through four built-in evaluation procedures that compare a detector’s outputs to a ground truth (GT) series of labels or events (see *Evaluation Procedures* section). GT labels may be derived from known aspects of the stimulus – such as the location and timing of external events (e.g., Komogortsev et al., 2010) – or manually-labeled data, as used in this report.

**Figure 1:**
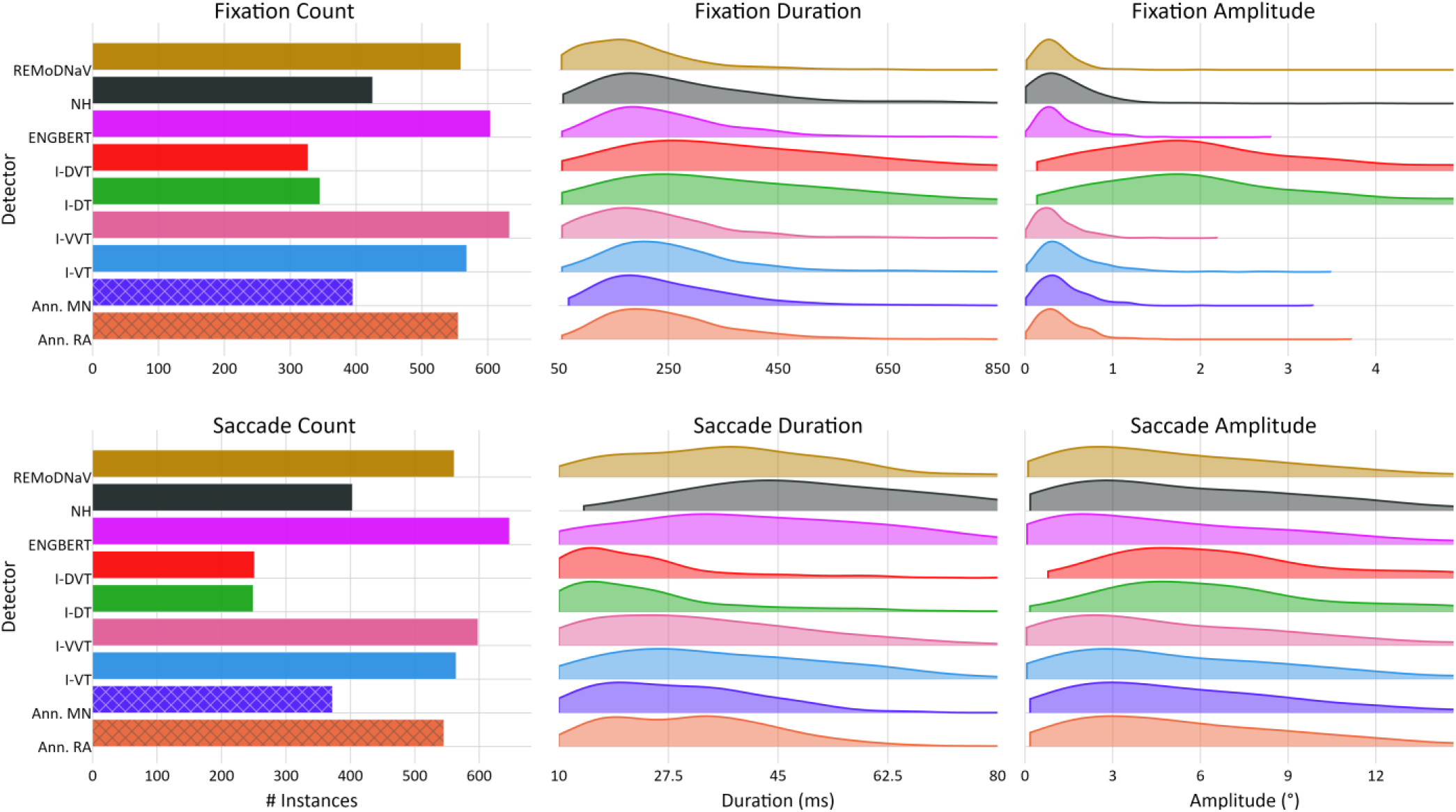
Distribution of EM Features Across Detectors and Annotators. *Note:* Comparison of fixation (top row) and saccade (bottom row) feature distributions for the *lund2013^+^-image* dataset, detected by human annotators (*RA* and *MN*) and algorithmic detectors implemented in *pEYES*. Shown features include the number of events (left), event duration (in milliseconds; middle), and event amplitude (in degrees visual angle; right).

To promote standardized evaluations, *pEYES* includes an API for downloading and parsing four publicly available ET datasets that include human-annotated data, which could be used as GT (Andersson et al., 2017; Hooge et al., 2018; Startsev et al., 2019; Zemblys et al., 2018). Descriptive statistics and metadata for these datasets are available in Table 1 of the review by Stratsev and Zemblys (2023). To evaluate detection algorithms for a specific use case, researchers can use *pEYES* to select a dataset that shares their data’s characteristics, such as stimulus type, sampling rate or recording duration.

We demonstrate how to implement a basic EM detection pipeline and visualize its output using *pEYES* in **Appendix B**. For a comprehensive overview of the package’s functionality, refer to the user manuals available at https://github.com/huji-hcnl/pEYES/tree/main/docs. The *pEYES* codebase, along with all analyses performed in this report, is available at: https://github.com/huji-hcnl/pEYES.

### Evaluation Procedures

Stratsev & Zemblys (2023) propose a taxonomy that distinguishes among three approaches for comparing and evaluating EM detection algorithms, each characterized by distinct metrics and methodologies: *Sample-Level Evaluation*, *Feature-Distribution Evaluation*, and *Event-Matching Evaluation*. The simplest evaluation procedure to implement is *Sample-Level Evaluation*, which compares the detector’s classification with the GT on a sample-by-sample basis. The inherent one-to-one mapping between sequences allows various metrics to be used to quantify agreement between detector and GT. For example, Andersson et al. (2017) used *Cohen’s Kappa* to quantify detectors’ performance, whereas Birawo and Kasprowski (2022) calculated *recall*, *precision* and *f1* scores for each event-type separately. Although relatively simple to implement, Startsev & Zemblys (2023) note that this approach may produce overly optimistic results due to the imbalanced nature of EM data. For instance, fixations often span many more data samples than saccades, potentially skewing metrics that do not account for this imbalance.

An alternative procedure, *Feature-Distribution Evaluation*, operates on the sets of EM *events* rather than the series of sample labels. This method compares the distribution of features – such as counts, duration, and amplitudes – between detected and GT events. For example, Andersson et al. (2017) calculated counts, means and standard deviations of durations separately for fixations, saccades and PSOs to quantify performance of 10 detectors with respect to human-annotated events. Similarly, Hooge et al. (2018) used event counts, durations, and velocities to examine the classification criteria of 12 human annotators. This type of evaluation procedure may provide “under the hood” insights into the classification process used by human or algorithm labelers, which may be helpful for parameter calibration. However, it has the inherent drawback of ignoring the sequential nature of the data, as events are aggregated and reduced to a low-dimensional (typically one- or two-dimensional) distribution.

Lastly, *Event-Matching Evaluation* procedures generate a mapping between detected and GT events and quantify match quality, either by calculating overall metrics like *recall* and *precision*, or by comparing features of the matched-events. For each detected event, the procedure searches for a corresponding event in the GT; if a match is found, various features of the matched pair – such as their duration difference or pupil size ratio – can be analyzed. Such procedures thus assess how well the detected events correspond to GT events, without requiring a sample-by-sample match as the Sample-Level Evaluation procedures. For instance, Hooge at al. (2018) introduced *Relative Timing Offset* (RTO) and *Relative Timing Deviation* (RTD) – the mean and standard deviation of paired events’ onset (or offset) differences – to assess the temporal alignment between detected and GT events. Alternatively, Kothari et al. (2020) used the 𝑙_2_*-distance of event-timings* and *Intersection-over-Union* (IoU) to measure the temporal alignment of matched events. This type of evaluation procedure strongly depends on user choices of matching schemes and parameters, which can produce varying mappings for the same GT and detected events.

These evaluation procedures offer complementary insights into detector performance, but are typically correlated, as they all depend on the same underlying factor: the detector’s ability to accurately align event boundaries – onsets and offsets – with the GT. A detector that precisely identifies the start and end of each event and labels intermediate samples accordingly, will produce a label sequence identical to the GT, thereby achieving maximal scores across all evaluation procedures and metrics. Consequently, detector performance assessment can be largely reduced to evaluating the detector’s sensitivity to event boundaries.

While the sensitivity index (𝑑′; Hautus et al., 2021) can be computed over the entire dataset to yield a single summary metric per event boundary (e.g., saccade onset), this approach does not support statistical comparisons between detectors. To enable such comparisons, we calculated 𝑑′ scores separately for eachrecording, allowing analysis of variability across detectors and event types.

To calculate sensitivity indices, we adapted the method described in Keren, Yuval-Greenberg & Deouell (2010, *Detection* section). An algorithm-detected event boundary (e.g., saccade onset) was considered a *hit* if a corresponding GT event boundary existed within a user-defined temporal window (±Δ𝑡). Unmatched GT and detected event boundaries were classified as *misses* and *false-alarms* (FAs), respectively. Based on these classifications, we computed the *true-positive rate* (TPr; i.e., *hit-rate*, *recall*), *positive predictive value* (PPV; i.e., *precision*), and the temporal-alignment metrics introduced above (e.g., *RTO*, *RTD*, and *temporal* 𝑙_2_*-distance*). We also defined the number of *negative windows* in the GT as the number of non-overlapping windows of size 2Δ𝑡 + 1 that contained no event boundary. This enabled calculating the detector’s *false-alarm rate* (FAr) and, together with the TPr, the sensitivity index (𝑑^′^ = 𝑧(𝑇𝑃𝑟) − 𝑧(𝐹𝐴𝑟))^2^.

### Dataset

Our analysis uses the extended *lund2013* dataset, originally published by Andersson et al. (2017). The original dataset included 34 recordings from 17 subjects; however, additional data was made public by the authors. This extended dataset, which we refer to as *“lund2013^+^”*, includes 63 recordings from 30 subjects, totalling 383,212 samples. Binocular gaze data was collected at a 500𝐻𝑧 sampling rate^3^ using a Hi-Speed1250 eye-tracker (SensoMotoric Instruments GmbH, Teltow, Germany), though only right-eye data is provided for analysis. To minimize head movement, subjects used a chin and forehead rest during recording.

Subjects were presented with one of three types of stimuli: a color image (87,790 samples), a real-world video (274,096 samples), or a linearly moving dot (21,326 samples). Two human annotators, *RA* and *MN*, labeled each sample in the dataset. Annotator *RA* labeled nearly all recordings (381,886 samples), while *MN* labeled only 34 files (104,745 samples). **Table 2** displays the proportion of samples assigned to each label by the annotators, illustrating the distribution of EM types within the dataset.

**Table 2:** Distribution of Eye Movement in the Extended lund2013^+^ Dataset.

| Stimulus | Annotator | Subjects | Sessions | # Annotated |  | % Labels |  |  |  |  |  | % Events |  |  |  |  |
| --- | --- | --- | --- | --- | --- | --- | --- | --- | --- | --- | --- | --- | --- | --- | --- | --- |
|  |  |  |  | Samples | Events | U/I | FIX | SAC | PSO | SP | BLI | FIX | SAC | PSO | SP | BLI |
| All | RA | 30 | 62 | 381886 | 5327 | 0.1 | 42.4 | 5.3 | 3.0 | 46.8 | 2.1 | 30.7 | 33.7 | 27.4 | 6.5 | 1.7 |
|  | MN | 20 | 34 | 104745 | 6939 | 0.3 | 61.5 | 7.2 | 4.4 | 22.6 | 4.1 | 22.4 | 34.4 | 25.2 | 16.9 | 1.2 |
| Image | RA | 18 | 20 | 87790 | 1586 | 0.1 | 76.5 | 9.2 | 4.8 | 4.9 | 4.7 | 36.2 | 33.7 | 27.9 | 0.3 | 2.0 |
|  | MN | 13 | 14 | 63849 | 1118 | 0.2 | 79.6 | 8.6 | 5.2 | 0.9 | 5.5 | 35.5 | 34.8 | 26.4 | 1.9 | 1.5 |
| Video | RA | 18 | 19 | 274096 | 3495 | 0.1 | 33.6 | 4.4 | 2.6 | 57.9 | 1.4 | 23.1 | 33.4 | 27.7 | 14.6 | 1.1 |
|  | MN | 9 | 9 | 29029 | 350 | 0.1 | 43.0 | 5.2 | 3.4 | 46.4 | 2.0 | 17.1 | 34.1 | 25.2 | 22.4 | 1.2 |
| Moving Dot | RA | 19 | 23 | 20000 | 246 | 1.0 | 12.8 | 4.7 | 1.4 | 79.5 | 0.5 | 6.9 | 34.0 | 22.9 | 35.4 | 0.7 |
|  | MN | 10 | 11 | 11867 | 144 | 1.4 | 9.0 | 4.5 | 2.0 | 81.6 | 1.4 | 13.0 | 34.6 | 16.7 | 35.4 | 0.4 |

*Note:* Percentage of samples and events assigned to each eye movement label – unidentified (U/I), fixation (FIX), saccade (SAC), post-saccadic oscillation (PSO), smooth pursuit (SP), and blink (BLI) – across three stimulus types (image, video, and moving dot), as annotated by the human annotators of the *lund2013<sup>+</sup>* dataset (RA and MN). Stimulus "All" represents aggregate statistics across all stimulus types. This breakdown highlights labeling patterns and differences between annotators at both the sample and event levels.

In this work, we focus on detection of fixations and saccades during static displays. Therefore, our analysis focuses only on *lund2013^+^-image*, the image-stimulus subset of the extended *lund2013^+^* dataset. This 87,790 samples subset was recorded from 18 different subjects in 20 sessions. Of these, 14 sessions (63849 samples) were part of the original dataset and were annotated by both *MN* and *RA*, and the remaining 6 sessions were annotated only by the latter. As shown in **Table 2**, fixations make up most of this data, accounting for 76.5% and 79.6% of samples labeled by *RA* and *MN*, respectively. Saccades are the second most frequent label, representing nearly 10% of the labels for both annotators. Notably, the annotators were unaware of the presented stimulus when labelling the data, leading to a small portion of samples being labeled as smooth pursuits (4.9% of *RA*’s labels), even though this type of EM is physiologically implausible when viewing a static stimulus.

### Detection Algorithms

We evaluate the performance of seven threshold-based detection algorithms, implemented within the pEYES package, as listed in **Table 1**. *I-VT* (Salvucci & Goldberg, 2000) calculates the sample-to-sample velocity of all data points and labels a sample as a saccade if its velocity exceeds a predefined threshold 𝑣_𝑠𝑎𝑐_, and as a fixation otherwise. The *I-VVT* algorithm (Komogortsev & Karpov, 2013) uses an additional velocity threshold, 𝑣_𝑠𝑝_, allowing it to detect fixations, saccades, and smooth pursuits that have an intermediate speed.

The *I-DT* algorithm relies on spatial dispersion to distinguish fixations from saccades. The implementation provided with pEYES follows the original algorithm proposed by Salvucci and Goldberg (2000), though different variations have been proposed over the years (Blignaut, 2009). In the original implementation, the sum of horizontal and vertical dispersion of gaze points within a predefined timeframe is compared with a predefined spatial threshold. If the dispersion is below the threshold, the window is extended to include additional samples, and the dispersion is recalculated. This process is repeated until the dispersion exceeds the threshold, then all samples in the window are labeled as a fixation, except the last sample which is labeled a saccade, and a new window begins from the subsequent sample.

Combining both *I-DT* and *I-VT*, the *I-DVT* algorithm (Komogortsev & Karpov, 2013) detects fixations using a spatial dispersion-threshold method. Then, in the remaining samples, it differentiates saccades from smooth pursuits using a velocity-threshold.

Originally developed to identify microsaccades within a fixation, *Engbert’s* algorithm (Engbert & Kliegl, 2003; Engbert & Mergenthaler, 2006) is commonly used to differentiate fixations from saccades in many ET experiments. Like I-VT, it uses a velocity threshold to determine whether a sample belongs to a fixation or a saccade, but unlike I-VT, this threshold is calculated individually for each recording session, based on the median-based standard deviation of velocities along the x-and y-axes.

The *NH* algorithm was developed by Nyström & Holmqvist (2010) to detect PSOs, fixations and saccades. Like *Engbert’s* algorithm, it calculates an adaptive velocity threshold, and applies it to detect saccade peak velocities. Once it detects samples with peak velocity, it traverses backwards and forwards in time to identify the saccade’s onset and offset, and the preceding PSO, if it exists.

Lastly, the *REMoDNaV* algorithm (Dar et al., 2021) builds on the *NH* algorithm, expanding it to support also the classification of smooth pursuits, making it applicable to both static and dynamic stimuli. Unlike *NH*, *REMoDNaV* segments the original data into chunks before calculating a locally adaptive velocity-threshold, based on robust statistics like the median absolute deviation. This approach is suggested to enhance robustness to noise and improve effectiveness with prolonged recordings, making *REMoDNaV* suitable for both high- and low-quality data.

### Detection Parameters

The *pEYES* package provides default values for detector parameters based on previous publications recommending optimal or default settings. However, in this report, where applicable, we use the parameter values proposed by Andersson et al. (2017) – minimum fixation duration (55𝑚𝑠), maximum fixation dispersion (2.7°), and maximum fixation velocity (45°/𝑠) – as these parameters were calibrated to match the decision criteria of human annotators in the original *lund2013* dataset. Additionally, we set a minimum duration for all EM types of 2 samples, as any single-sample event at a 500𝐻𝑧 sampling rate is likely to represent an erroneous label. For saccades specifically, we require a minimum duration of 10𝑚𝑠, which is the default value used by the *NH* (Nyström & Holmqvist, 2010) and *REMoDNaV* (Dar et al., 2021) algorithms. Lastly, we use an intermediate velocity threshold of 26°/𝑠 for the *I-VVT* detector to distinguish fixations from smooth pursuits, as proposed in the original description of this algorithm by Komogortsev and Karpov (2013). All other parameters are set to their default values. A comprehensive list of parameter and argument values used in this analysis is available in **Appendix C**.

To assess how well these parameter values apply to the *lund2013^+^-image* dataset, we calculated the proportion of human-annotated events that satisfy each criterion (excluding single-sample events and smooth pursuits, see **Table 3**). Over 90% of annotated events meet the predefined duration thresholds. The dispersion threshold (2.7°) covers over 98% of fixations and 79% of saccades, meaning 21% of saccades fall below this spatial threshold – potentially hindering performance of spatially-tuned detectors. While over 99% of saccades exceed the velocity threshold (45°/𝑠), most (over 75%) have a *minimum* velocity below it, which may impair velocity-based algorithms’ ability to precisely detect saccade onsets and offsets. Additionally, 28% of fixations exceed this threshold, risking their misclassification as saccades by such velocity-based detectors. Although these thresholds do not perfectly fit the dataset, we apply them in our work to maintain consistency with the analysis performed by Andersson et al., thereby facilitating comparability with their findings. Interested readers can use the open source *pEYES* to test other parameters.

**Table 3:**
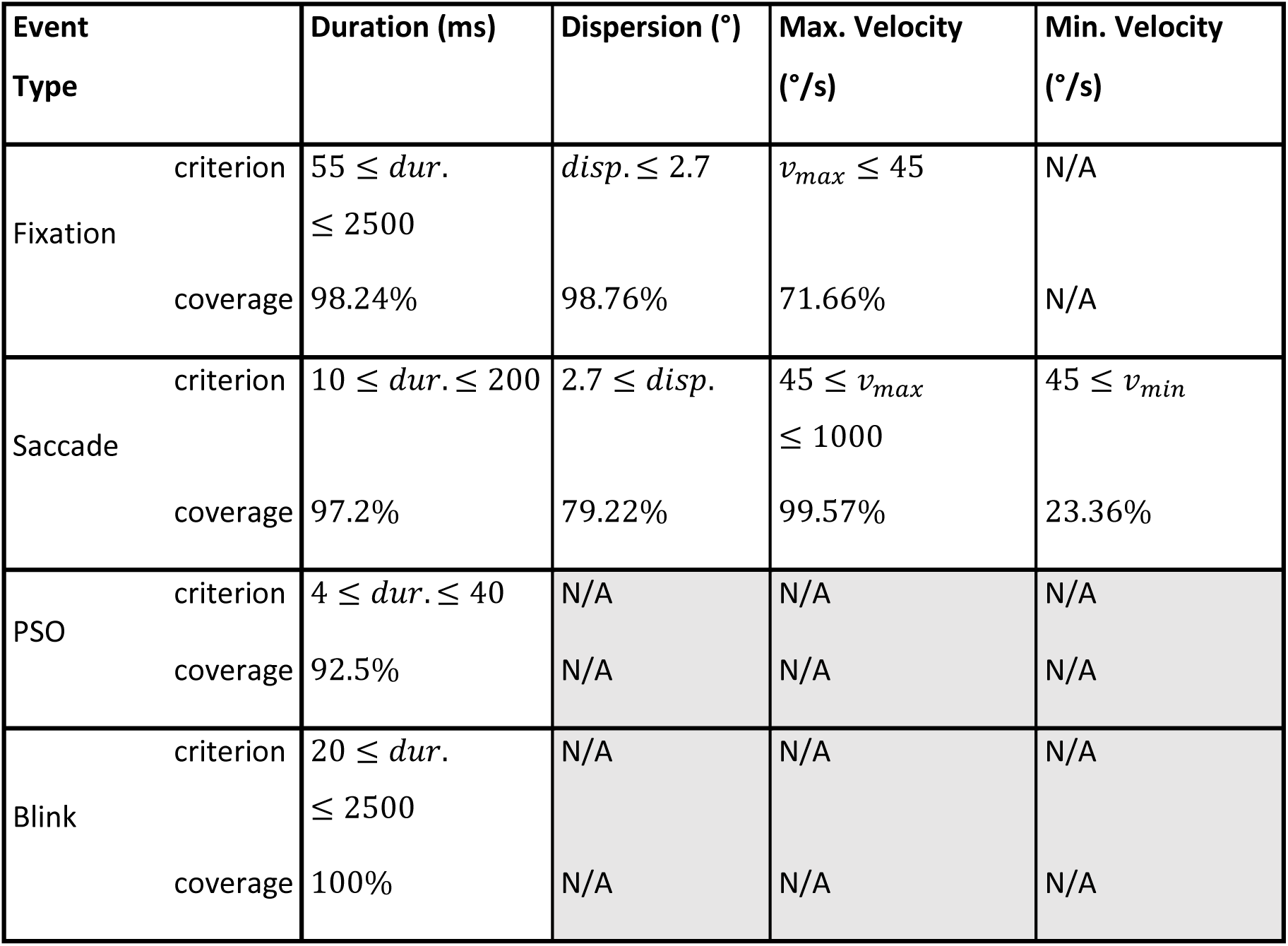
Percent of Ground-Truth Events Satisfying the Default Thresholds.

## Results

### Sample-Level Evaluation

#### Sample-by-Sample Agreement

We started by evaluating detector performance across all event types, by measuring the sample-by-sample agreement between the detected and GT labels for each recording. Following Andersson et al., we measured agreement using *Cohen’s Kappa* (Cohen, 1960), as shown in the top panel of **Figure 2**. We conducted a Friedman’s test (Friedman, 1937) comparing detectors’ *Kappa* scores for each annotator separately, and found a significant difference for both annotator *RA* (𝑄^𝑅𝐴^(6) = 76.5, 𝑝 < 0.001) and annotator *MN* (𝑄^𝑀𝑁^(6) = 61.6, 𝑝 < 0.001). Pairwise post-hoc comparisons using the Wilcoxon-Nemenyi-McDonald-Thompson test (also known as Tukey-HSD; Hollander et al., 2013; Pereira et al., 2015) revealed three significantly distinct subsets of detectors. The top-ranking subset, consisting of *I-VT*, *I-VVT*, and *Engbert*, significantly outperformed the bottom-ranking subset, which includes *I-DT* and *I-DVT*, and (all 𝑝_𝑝𝑎𝑖𝑟𝑤𝑖𝑠𝑒_ ≤ 0.05; note the Tukey-HSD test incorporates a correction for multiple comparisons). The *NH* and *REMoDNaV* detectors were classified within a middle-ranking subset, which did not differ significantly from either the top or bottom subsets.

**Figure 2:**
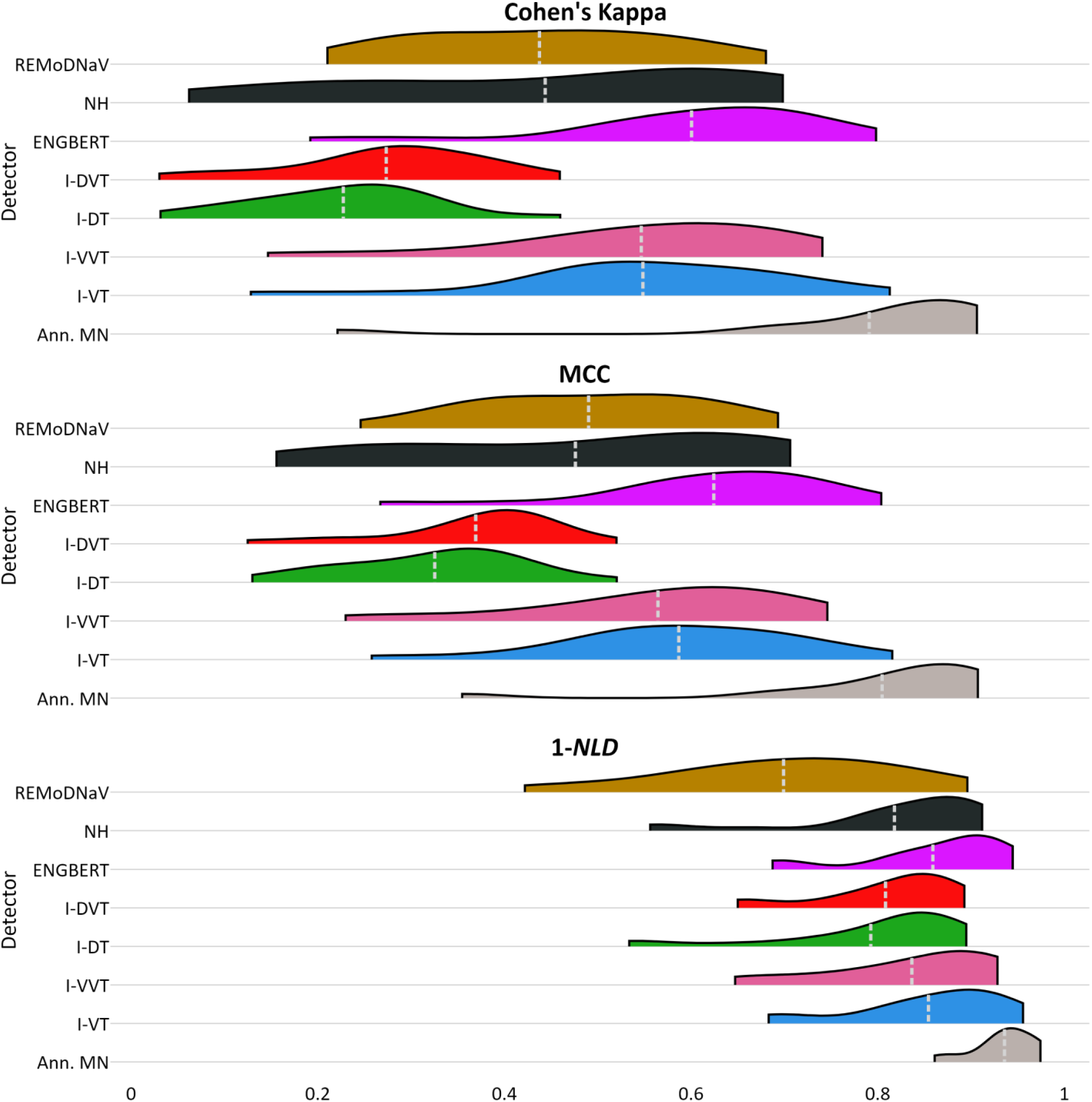
Distribution of Sample-Level Agreement Scores Between Annotator RA and Algorithmic Detectors. *Note:* Distribution of sample-by-sample agreement scores (*Cohen’s Kappa*, *MCC*, *1-NLD*) between GT annotator *RA* and each of the algorithmic detectors implemented in *pEYES*, across recordings. Inter-rater agreement (with annotator *MN*) is provided for comparison. The mean of each distribution is marked by a dashed light-gray line. A similar figure based on *MN* as ground truth is provided in **Appendix D**.

We repeated this analysis using Matthews Correlation Coefficient (*MCC*; middle panel of **Figure 2**) and Complementary Normalized Levenshtein Distance (*1-NLD*; bottom panel of **Figure 2**), introduced by Startsev and Zemblys (2023). MCC scores produced a similar pattern, whereas *1-NLD* was less sensitive to performance differences, consistent with previous finding suggesting this metric to be overly-optimistic (Startsev & Zemblys, 2023). Statistical analyses of *MCC* and *1-NLD*, along with pairwise comparison results, are available in **Appendix D**.

#### Fixation and Saccade Sensitivity

To further assess sample-level performance, we calculated each detector’s sensitivity indices (𝑑′) for fixations and saccades. A sample was considered a *hit* if it matched the corresponding label in the GT, or a *false-alarm* (FA) otherwise. Likewise, GT-labeled fixation (or saccade) samples that were mis-labeled by the detector were categorized as *misses*, and all remaining samples were categorized as *correct rejections* (CRs). Using these categorizations, we computed the sensitivity index (𝑑′) for fixations (or saccades) for each detector in each recording, applying a log-linear correction for extreme proportions where required (Hautus, 1995).

Statistical comparisons were conducted separately using each human annotator (*RA* and *MN*) as ground truth. Friedman tests revealed significant differences in detector sensitivity for both fixations (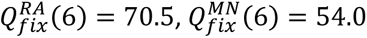; both 𝑝_𝑠_ < 0.001) and saccades (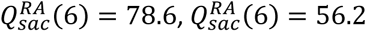; both 𝑝_𝑠_ < 0.001; superscript and subscript indices denote the annotator and event type, respectively). Post-hoc pairwise comparisons using a Tukey-HSD test indicated that *REMoDNaV* and *Engbert* performed significantly better in saccade sensitivity than *I-DT*, *I-DVT* and *Engbert’s* detector. The *I-VT* detector demonstrated the highest fixation sensitivity, significantly outperforming *REMoDNaV*, *NH*, and *I-DT* (for detailed results, see **Appendix E**).

#### Event Matching Evaluation

Sample-level evaluation may overestimate performance due to the dominance of fixation samples in ET data (Startsev & Zemblys, 2023), or underestimate it in cases where precise onset and offset detection is difficult to achieve. As a complementary approach, we evaluated detection performance based on fixation and saccade matching, using both *temporal alignment* measures and *boundary sensitivity* indices to quantify performance. To that end, we paired detected event onsets and offsets with the nearest (smallest absolute temporal difference) comparable event in the GT data. A detected onset (or offset) was included in the analysis only if its corresponding GT onset (or offset) was found within a temporal window of |Δ𝑡| ≤ 20 𝑠𝑎𝑚𝑝𝑙𝑒𝑠^4^.

#### Temporal Alignment Measures

Figure 3 depicts the distribution of temporal differences among successfully matched fixation onsets and offsets (top- and bottom-left, respectively), and saccade onsets and offsets (top- and bottom-right, respectively), using *RA*’s annotations as GT (for *MN* as GT, yielding similar results, see **appendix F**). For each distribution, the mean and standard deviation represent the detector’s Relative Temporal Offset and Relative Temporal Deviation, respectively (RTO and RTD; summarized in **Table 4**). Unmatched onsets and offsets – falling outside the temporal window – were excluded from these calculations and subsequent analyses.

**Figure 3:**
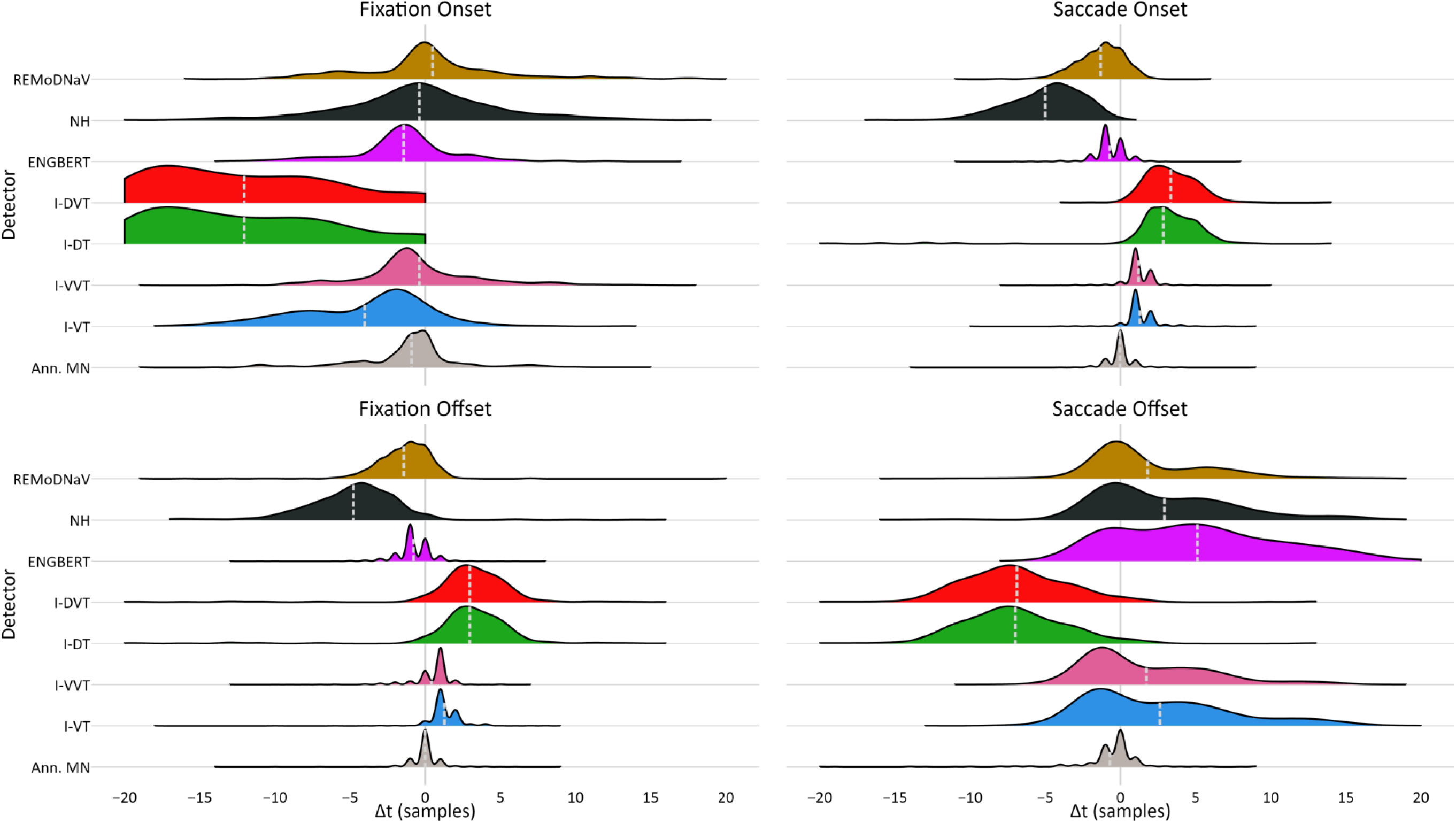
Distribution of Temporal Misalignments Relative to Fixation and Saccade Boundaries Identified by RA. *Note:* Distribution of temporal alignments (in samples) between detected fixation and saccade onsets and offsets that were matched to corresponding events annotated by *RA*. Alignment values were truncated to the range [-20, 20] 𝑠𝑎𝑚𝑝𝑙𝑒𝑠. The mean of each distribution, marked by a dashed light-gray line, is the detector’s *RTO*, and the distribution’s spread (i.e., its standard deviation) reflects its *RTD*. Both values are specified in **Table 4**. A similar figure based on *MN* as ground truth is provided in **Appendix F**.

**Table 4:** Temporal Alignment of Fixation and Saccade Boundaries. *Note:* Hit-rate (in %), RTO, and RTD (both in sample units**^Error!^ ^Bookmark^ ^not^ ^defined.^**) for matches between detected and human-annotated fixation and saccade onsets and offsets. For each GT annotator, the top (highlighted) row shows alignment scores of the 2^nd^ annotator. Within each column, the two highest performing detectors are shown in **bold**.

| GT | Detector | Fixation Onset |  |  | Fixation Offset |  |  | Saccade Onset |  |  | Saccade Offset |  |  |
| --- | --- | --- | --- | --- | --- | --- | --- | --- | --- | --- | --- | --- | --- |
|  |  | Hit-Rate | RTO | RTD | Hit-Rate | RTO | RTD | Hit-Rate | RTO | RTD | Hit-Rate | RTO | RTD |
| RA | MN | 95.9% | -0.9 | 4.0 | 95.2% | 0.0 | 1.4 | 98.7% | 0.0 | 1.2 | 98.4% | -0.7 | 2.9 |
|  | I-VT | 92.2% | -4.0 | 4.4 | 93.8% | -1.3 | **1.5** | 93.7% | 1.3 | **1.2** | 93.1% | 2.6 | 5.0 |
|  | I-VVT | **97.5%** | **-0.4** | **4.2** | **98.0%** | **0.4** | 1.6 | 96.0% | **1.2** | **1.2** | 95.5% | **1.7** | 4.3 |
|  | I-DT | 39.3% | -12.1 | 5.8 | 55.6% | 3.0 | 3.7 | 52.2% | 2.8 | 3.9 | 52.0% | -7.0 | **3.6** |
|  | I-DVT | 39.3% | -12.1 | 5.8 | 55.6% | 3.0 | 3.7 | 52.9% | 3.3 | 1.7 | 52.7% | -6.9 | **3.9** |
|  | Engbert | **98.0%** | -1.4 | **3.7** | **99.0%** | **-0.8** | **1.4** | **99.1%** | **-0.7** | **1.2** | **98.7%** | 5.1 | 5.4 |
|  | NH | 75.0% | **-0.3** | 5.9 | 76.9% | -4.9 | 3.4 | 76.8% | -5.1 | 2.7 | 75.2% | 3.1 | 5.1 |
|  | REMoDNaV | 86.1% | 0.5 | 5.2 | 72.6% | -1.4 | 2.4 | **98.2%** | -1.3 | 1.6 | **98.0%** | **1.8** | 4.5 |
| MN | RA | 92.8% | 0.9 | 4.0 | 95.0% | 0.0 | 1.4 | 97.9% | 0.0 | 1.2 | 97.6% | 0.7 | 2.9 |
|  | I-VT | 94.1% | -3.8 | 4.8 | 95.3% | 1.2 | **1.4** | 95.0% | 1.3 | **1.3** | 93.6% | 3.4 | 5.7 |
|  | I-VVT | **97.8%** | **-0.1** | **4.3** | **98.3%** | **0.3** | 1.6 | **97.6%** | **1.2** | **1.3** | 96.6% | **2.1** | 5.3 |
|  | I-DT | 43.3% | -12.9 | 6.3 | 55.9% | 3.0 | 3.2 | 52.3% | 2.9 | 3.6 | 52.5% | -6.2 | **4.5** |
|  | I-DVT | 43.1% | -13.0 | 6.1 | 55.9% | 3.0 | 3.2 | 53.1% | 3.4 | 2.0 | 53.1% | -6.1 | **4.5** |
|  | Engbert | **99.0%** | -1.3 | **4.1** | **99.2%** | **-0.7** | **1.5** | **98.9%** | **-0.7** | **1.2** | **98.4%** | 6.1 | 5.7 |
|  | NH | 72.5% | **-0.2** | 5.8 | 76.0% | -5.0 | 3.8 | 74.0% | -5.3 | 2.8 | 72.9% | 3.7 | 4.8 |

#### Fixation Temporal Alignment

The two human annotators demonstrated high agreement in fixation timing, with over 95% of RA’s annotated fixations matching MN’s (both for onsets and offsets). The RTO and RTD for fixation onsets were 0.9 and 4.0 samples, respectively; and for offsets 0.0 and 1.4 samples, respectively. Notably, both *I-VVT* and *Engbert’s detector* outperformed or matched this inter-annotator alignment, achieving hit-rates of 97.5% and *RTO-RTD* values comparable to or better than those of the second annotator.

We conducted a Kruskal-Wallis test (Kruskal & Wallis, 1952) to compare temporal differences across detectors for each annotator. Significant differences were found for both fixation onsets 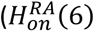 = 997.7, 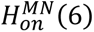 = 751.9; both *p_s_* < 0.001) and offsets (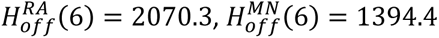; both *p_s_* < 0.001). A post-hoc pairwise Dunn’s test applied with a Bonferroni correction for multiple-comparisons (Abdi, 2007; Bonferroni, 1936; Dunn, 1964), revealed significant differences between most detector pairs, regardless of GT annotator and boundary type (onset or offset; see **Appendix F3 and F4**)^5^.

#### Saccade Temporal Alignment

Saccade timing was also highly consistent between annotators, with an onset hit-rate of 98% and RTO and RTD measures of 0.0 and 1.2 samples, respectively. Similarly, nearly 98% of saccade offsets were successfully matched, with an RTO of 0.7 and RTD of 2.9 samples. The top performing algorithms were, again, *I-VVT* and *Engbert’s* detector, as well as *REMoDNaV*. All three achieved comparable alignment scores, closely approaching the performance of the second human annotator.

As with fixations, a Kruskal-Wallis test revealed significant differences across detectors for both saccade onset (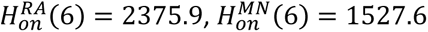; both 𝑝_𝑠_ < 0.001) and offset (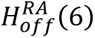 = 1301.5, 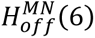 = 793.3; both 𝑝_𝑠_ < 0.001). Post-hoc Dunn’s test with a Bonferroni correction revealed significant differences between most detector pairs, regardless of GT annotator and boundary type (see **Appendix F5 and F6**)^5^.

#### Boundary-Sensitivity Evaluation

Temporal alignment measures offer valuable insights into detector performance, but interpreting them often requires evaluating multiple measures simultaneously (e.g., RTO, RTD, and hit-rate). In contrast, *event-boundary sensitivity evaluation* relies on a single metric, the sensitivity index (𝑑^′^). We applied incremental temporal windows (|Δ𝑡| ≤ 0,1, …,20 𝑠𝑎𝑚𝑝𝑙𝑒𝑠) to categorize event boundaries (onsets or offsets) as *hits*, *misses*, and *FAs*. Hit rates were calculated as the number of matched algorithm-detected events divided by the number events in the GT data. False alarm rates were calculated as the number of unmatched algorithm-detected events divided by the *negative count*. The *negative count* was defined as the number of non-overlapping time-windows of size 2Δt + 1 in the GT data that do not contain an event boundary. Using these classifications, we computed each detector’s onset (or offset) sensitivity index (𝑑′) for each recording, applying a log-linear correction in cases where no hits or no false alarms were detected (Hautus, 1995).

Figures 4 **and 5** depict detectors’ sensitivity index scores (𝑑′) for fixations and saccades, respectively, using annotator *RA* as GT: the top row shows sensitivity scores for increasing Δ𝑡 values, and the bottom row shows the distribution of 𝑑′ scores across recordings for a strict temporal threshold of |Δt| ≤ 5 𝑠𝑎𝑚𝑝𝑙𝑒𝑠 (equivalent to 10𝑚𝑠 for most available recordings^4^).These results indicate that, across most temporal thresholds, fixation onsets and offsets were best detected by *Engbert’s* detector; saccade onsets were most accurately identified by both *REMoDNaV* and *Engbert’s* detector; and *REMoDNaV* showed the highest sensitivity to saccade offsets.

**Figure 4:**
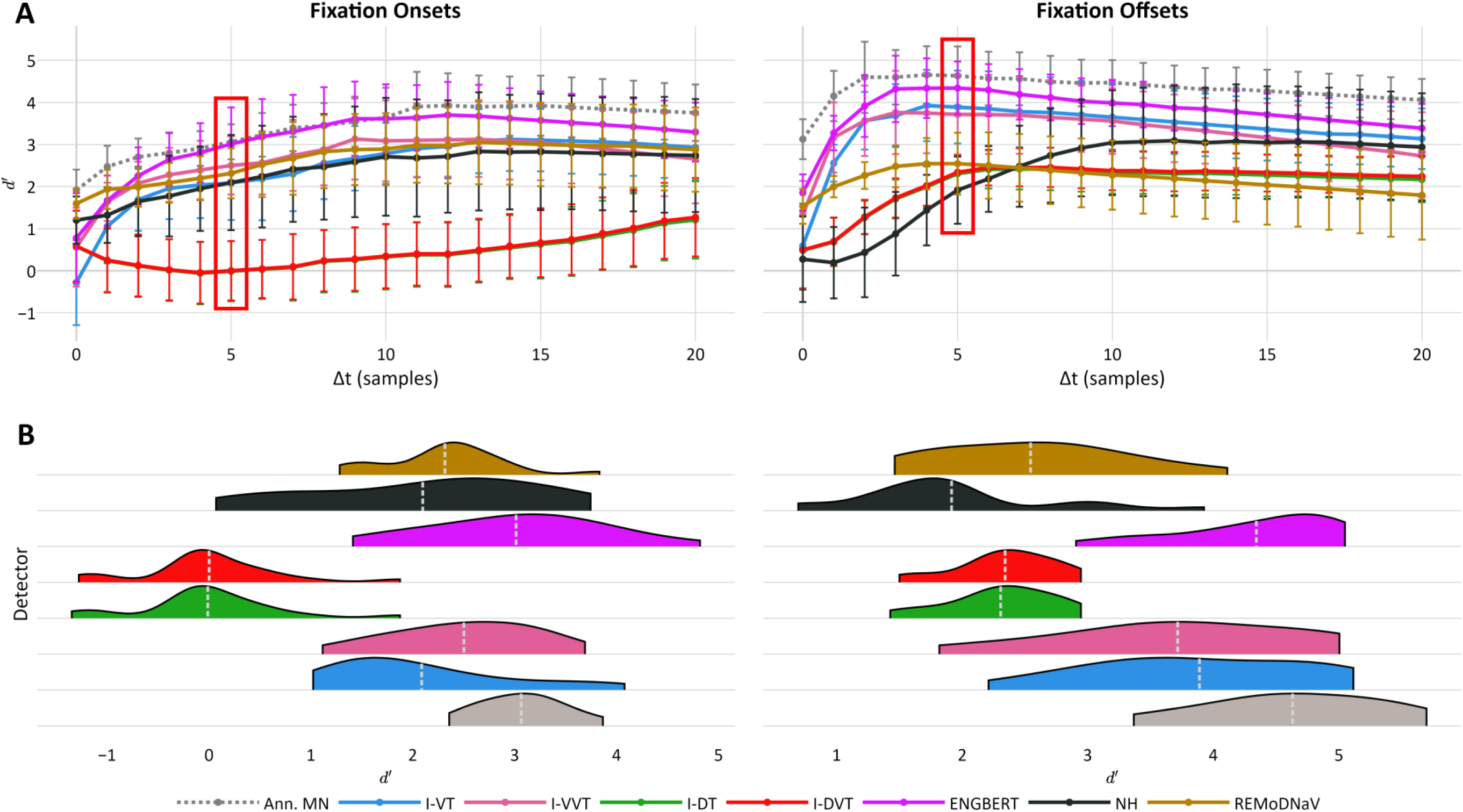
Fixation Boundary Sensitivity Index (d′) Across Temporal Thresholds Relative to Annotator RA. *Note:* Sensitivity index (𝑑′) scores for fixation onset and offset detection by each detector, using human annotator *RA* as GT. Sensitivity scores of the second human annotator (*MN*) are shown for reference (gray line and violin). (**A**) Mean sensitivity scores across increasing temporal windows (|Δ𝑡| = 0,1, …,20 𝑠𝑎𝑚𝑝𝑙𝑒𝑠). Each line corresponds to a detector’s mean 𝑑′ across recordings, with error bars indicating standard deviation. The red box marks the Δ𝑡 = 5 threshold, depicted in detail in Panel B. (**B**) Distribution of 𝑑′ scores at the selected Δ𝑡 = 5 threshold. White dashed lines mark the mean 𝑑′ score of each distribution. A similar figure using *MN* as ground truth is provided in **Appendix G**.

**Figure 5:**
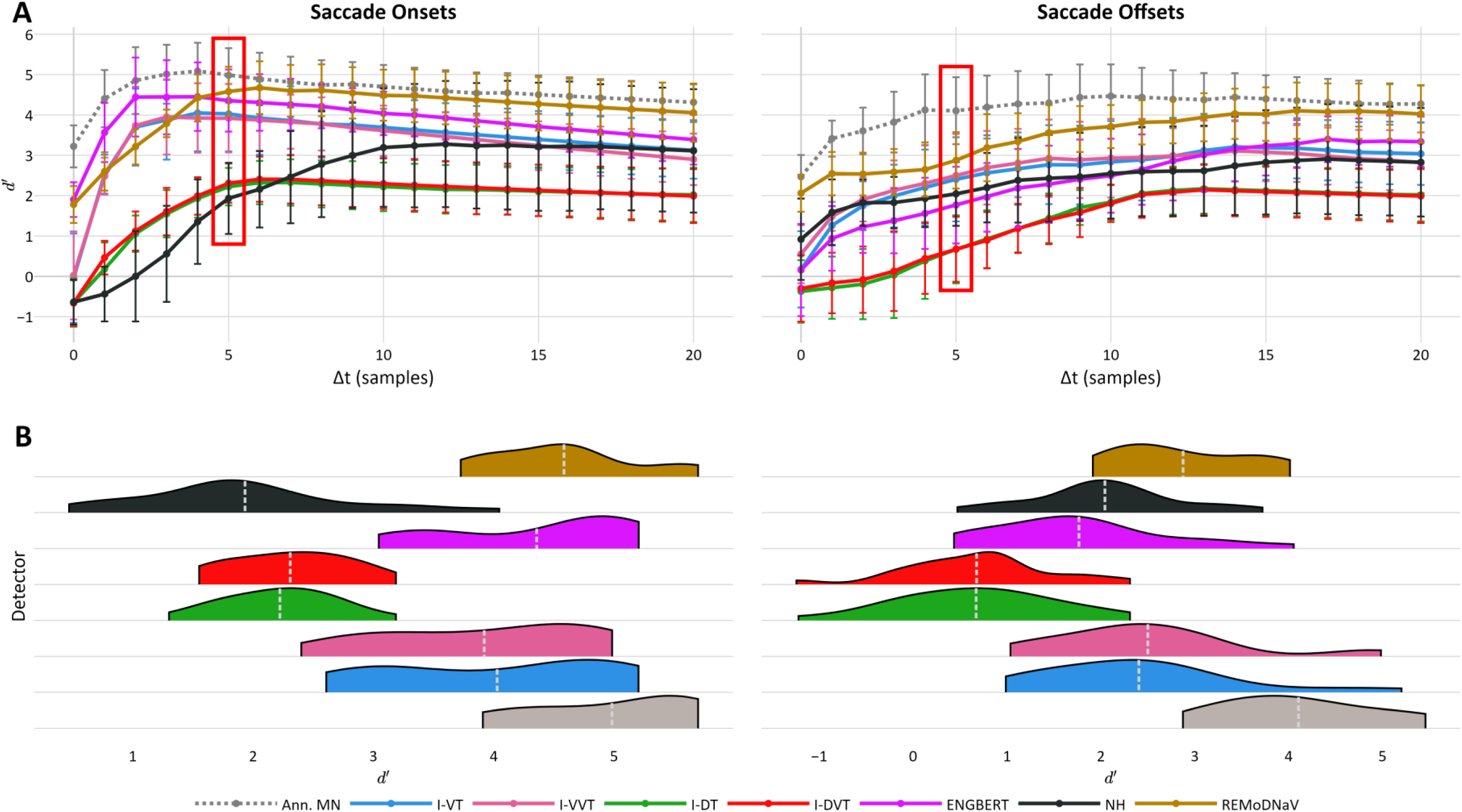
Saccade Boundary Sensitivity Index (d′) Across Temporal Thresholds Relative to Annotator RA. *Note:* Sensitivity index (𝑑′) scores for saccade onset and offset detection by each detector, using human annotator *RA* as GT. Sensitivity scores of the second human annotator (*MN*) are shown for reference (gray line & violin). (**A**) Mean sensitivity scores across increasing temporal windows (|Δ𝑡| = 0,1, …,20 𝑠𝑎𝑚𝑝𝑙𝑒𝑠). Each line corresponds to a detector’s mean 𝑑′ across recordings, with error bars indicating standard deviation. The red box marks the Δ𝑡 = 5 threshold, depicted in detail in panel B. (B) Distribution of 𝑑′ scores at the selected Δ𝑡 = 5 threshold. White dashed lines mark the mean 𝑑′ score of each distribution. A similar figure using *MN* as ground truth is provided in **Appendix H**.

These results indicate that, across most temporal thresholds, fixation onsets and offsets were best detected by *Engbert’s* detector; saccade onsets were most accurately identified by both *REMoDNaV* and *Engbert’s* detector; and *REMoDNaV* showed the highest sensitivity to saccade offsets. Conversely, detectors *I-DT* and *I-DVT* underperformed across nearly all tasks and thresholds. Notably, similar patterns emerge when using *MN* as the GT annotator (see **Appendices G and H**), and when applying the same analysis using 𝑓1 score as an alternative to 𝑑′ (results are available online).

We conducted Friedman tests to compare scores across detectors, while fixing the temporal threshold to Δ𝑡 ≤ 5 𝑠𝑎𝑚𝑝𝑙𝑒𝑠. These analyses revealed significant differences in detection sensitivity for fixation onsets 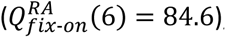, fixation offsets 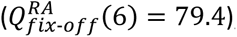, saccade onsets 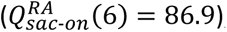, and saccade offsets 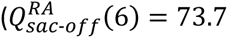; all 𝑝_𝑠_ < 0.001). Pairwise post-hoc comparisons using the Tukey-HSD test showed that *I-DT* and *I-DVT* significantly underperformed in fixation onset detection compared to all other detectors. Engbert’s detector consistently performed as well as or better than all other detectors for fixation onsets and offsets (𝑎𝑙𝑙 𝑝_𝑝𝑎𝑖𝑟𝑤𝑖𝑠𝑒_ ≤ 0.05; see **Appendix G3 and G4**). For saccade onsets, two distinct performance tiers emerged: *I-VT*, *I-VVT*,*Engbert* and *REMoDNaV* significantly outperformed *I-DT*, *I-DVT*, and *NH*. A similar pattern was observed for saccade offsets, where *REMoDNaV*, *NH*, *I-VT*, and *I-VVT* performed significantly better than *I-DT* and *I-DVT*. *Engbert’s* detector produced intermediate results: significantly lower than the top performing algorithm, *REMoDNaV*, but not significantly different than the remaining top- and bottom-ranking detectors (see **Appendix H3 and H4**).

### Validation Study: HFC-image Dataset

To validate our findings, we applied the same analysis steps to a different dataset, the Human Fixation Classification (*HFC*) dataset (Hooge et al., 2018). Unlike the *lund2013* dataset, the *HFC* dataset was recorded using a Tobii TX300 eye-tracker (Tobii Technology, Stockholm, Sweden) at a sampling rate of 300𝐻𝑧, providing alternative recording conditions for evaluation. Within this dataset we focused on the *HFC-image* subset, which consists of 10 recordings of free-viewing static color images, comprising 45,018 samples labeled by 12 human annotators as either “fixation” or “not fixation” (other event types were not identified). Annotators *RA* and *MN*, who also annotated the *lund2013* dataset, classified 82% and 84% of samples as fixations, identifying 753 and 750 distinct fixation events, respectively.

We applied the same seven detectors to the *HFC-image* dataset, using identical parameters as in the primary analysis, and evaluated their performance against *RA* and *MN*’s labels. Consistent with previous findings, the *Engbert* detector performed as well as or better than all other algorithms in sample-by-sample agreement with the GT, fixation temporal alignment, and fixation onset and offset sensitivity (see **Appendices I, J, and K**, respectively). However, contrary to the previous findings, dispersion-threshold detectors (*I-DT* and *I-DVT*) significantly outperformed velocity-threshold algorithms (*I-VT* and *I-VVT*) across most performance measures. Notably, these results were replicated when the analysis was repeated using GT annotations by other ET experts: *DN*, who is from the same lab as *RA* and *MN*, as well as *IH* and *JV*, who are affiliated with a different institution (example results are available online).

### Onset-Offset Comparison

Following our boundary-sensitivity results, we conducted an exploratory analysis comparing 𝑑′ scores for saccade onsets versus offsets across all detection algorithms, using the stringent Δ𝑡 ≤ 5 𝑠𝑎𝑚𝑝𝑙𝑒𝑠 temporal window for computing 𝑑′.We hypothesized that saccade onsets would be easier than offsets to detect, when compared across all recordings in the *lund2013^+^-image* dataset and all *pEYES’* detectors. This hypothesis was confirmed via a Wilcoxon Signed-Rank test for repeated measures (Wilcoxon, 1945), conducted using either annotator *RA* as GT (𝑊 = 9348, 𝑝 < 0.001), or annotator *MN* as GT (𝑊 = 4671, 𝑝 < 0.001; see Figure 6).

**Figure 6:**
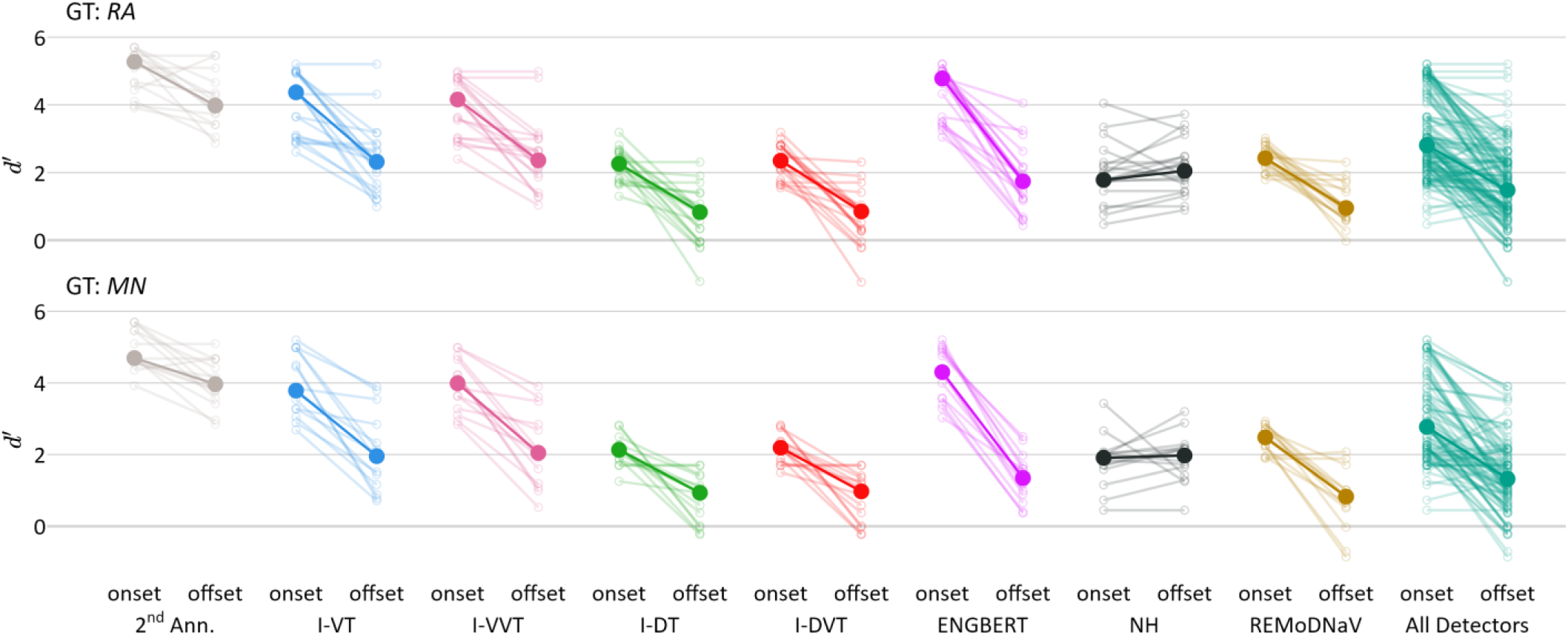
Detection Sensitivity Index (𝑑′) for Saccade Onsets and Offsets. *Note:* Saccade onset and offset sensitivity indices (𝑑′) for each detector, using human annotators *RA* (top row) and *MN* (bottom row) as GT, based on the *lund2013*^+-^*image* dataset. Sensitivity was computed using a temporal window of Δ𝑡≤5 𝑠𝑎𝑚𝑝𝑙𝑒𝑠. Small, light-colored circles represent individual recordings, with lines connecting onset and offset 𝑑′scores from the same recording. Large, opaque circles represent each detector’s median 𝑑′ score. The “All Detectors” column (in green-teal) reflects the overall distribution across all detectors. Results from the second annotator (2^nd^ Ann.) are shown for comparison (in light gray). A corresponding figure for fixation onset and offset detection sensitivity is provided in **Appendix L**.

We repeated this analysis for fixation onsets versus offsets and, as expected, found complementary results: fixation offsets were significantly easier to detect than onsets across all detectors, regardless of GT annotator (*RA* or *MN*) or dataset (*lund2013^+^-image* or *HFC-image*). Detailed results are reported in **Appendix L**.

## Discussion

We introduce ***pEYES***, an open-source Python toolkit designed to standardize EM detection and facilitate robust comparisons between detection algorithms. By applying this package to two publicly available datasets, we demonstrated how different detectors vary in their agreement with human annotations, their temporal alignment with event boundaries, and their sensitivity to fixation and saccade onsets and offsets. This evaluation highlights the performance of *Engbert*’s adaptive velocity-threshold algorithm (Engbert & Kliegl, 2003; Engbert & Mergenthaler, 2006), while also underscoring the impact of dataset characteristics and hyperparameter selection on detector performance.

The primary goal of this research was to address the variability and lack of transparency often associated with EM detection algorithms. We applied *pEYES* to two publicly available eye-tracking datasets – *lund2013^+^-image* and *HFC-image* – using human annotated labels as ground truth for comparing detection performance of seven commonly used threshold-based detectors. Detectors were evaluated based on multiple criteria, including **label agreement** with the GT (*Cohen’s Kappa*, *MCC*, *1-NLD*), **temporal alignment** of fixations and saccades (*hit-rate*, *RTO*, *RTD*), and **event boundary sensitivity** to fixation and saccade onsets and offsets (𝑑′). The latter is a novel metric we introduced for quantifying detection performance relative to a GT sequence. This metric requires specifying only a single parameter: the temporal window (±Δ𝑡) within which detected and GT event-boundaries are considered a match. By computing the detector’s sensitivity index (𝑑′) for each event-boundary (e.g., fixation onset, saccade offset), this method provides a concise, interpretable measure of performance. Importantly, it does not require detection of other types of events or boundaries, thereby making it suitable for analyses focused on a single class of events or for cases where only partial annotations are available.

Our findings revealed substantial differences in detector performance across datasets, evaluation metrics, event types, and event boundaries. Engbert’s adaptive velocity-threshold algorithm (Engbert & Kliegl, 2003; Engbert & Mergenthaler, 2006) consistently matched or outperformed all other detectors in both datasets, occasionally achieving human-level performance. The other adaptive-threshold algorithms – NH (Nyström & Holmqvist, 2010) and REMoDNaV (Dar et al., 2021) – performed comparably to Engbert’s in several tasks, with REMoDNaV narrowly outperforming it in saccade offset detection. These patterns remain consistent across datasets and GT annotators, underscoring the robustness of adaptive-threshold detectors.

Further analysis revealed that many of REMoDNaV’s errors stemmed from misclassifying fixations as smooth pursuits. Manually correcting these labels led to significant improvements in detector performance, both for fixation sensitivity and sample-by-sample agreement with the GT, at times matching Engbert’s performance (results are available online).

Unlike adaptive-threshold algorithms, global-threshold algorithms – such as I-VT, I-DT (Salvucci, D. D., & Goldberg, 2000), I-VVT, and I-DVT (Komogortsev & Karpov, 2013) – rely on predefined spatial, velocity, or acceleration thresholds to classify samples. While this makes them simple to implement and computationally efficient, their fixed thresholds reduce detection flexibility, hindering performance. This limitation was evident in our results: while these detectors performed well on one dataset, they generalized poorly when applied to the other. This result reinforces the benefits of adaptive-threshold algorithms.

Although Engbert’s detector performed consistently well across detection tasks and evaluation measures, all detectors exhibited performance variability depending on the context: the dataset, metric, event type, and boundary. This aligns with previous findings by Andersson et al. (2017), and is echoed in other evaluation domains. For example, Reich et al. (2025) recently reported substantial performance differences between I-VT and I-DT in psycholinguistic analysis and machine learning classification. Together, these findings emphasizes that **no single detector is universally optimal**, highlighting the importance of standardized, task-specific, detector-selection process, and demonstrate the practical value of incorporating *pEYES* into eye-tracking analysis pipelines.

### Event Onset and Offset Detection

Our analysis revealed that saccade onsets were easier to detect than offsets, and the opposite was true for fixations. This reduced sensitivity to the saccade-to-fixation transition was consistent across GT annotators and datasets, suggesting a shared limitation among all evaluated algorithms.

Importantly, accurate detection of this transition is critical in perceptual and cognitive research, where it is frequently used as a temporal anchor for analyzing neural responses during free viewing tasks. For instance, many studies examine fixation-related potentials (FRPs) – EEG signals time-locked to fixation onsets – to study neural responses during natural viewing and visual search tasks (e.g., Auerbach-Asch et al., 2020; Qiu et al., 2023). These analyses rely on precise determination of fixation onset, yet our findings suggest that achieving such temporal precision is more feasible for the complementary transition, from fixation to saccade onset. Notably, recent research indicates that neural responses during naturalistic viewing are more tightly aligned to saccade onset than to fixation onset (for example, Amme et al., 2024; Auerbach-Asch et al., 2023). Indeed, the top-performing algorithms identified in this report – Engbert’s detector and REMoDNaV – are well suited for analyses that rely on saccade-locked neural responses.

### Other Toolboxes

The ***pEYES*** package contributes new functionality to the growing ecosystem of Python tools for eye-tracking researchers. Other packages, such as *PyGaze* (Dalmaijer et al., 2014) and *PsychoPy* (Peirce et al., 2019), enable researchers to design and run experiments that incorporate eye tracking, while *PyTrack* (Ghose et al., 2020) supports statistical analysis of ET and EM data. More recently, *pymovements* (Krakowczyk et al., 2023) introduced standardized Python interfaces for working with multiple eye-tracking datasets, including implementations of three commonly used detectors.

While addressing similar practical needs, *pEYES* was developed to complement these tools by focusing on detector standardization and performance evaluation. It includes a broad set of well-documented detection algorithms, and offers a comprehensive framework for anaslyzing and visualizing detector performance across multiple metrics and evaluation procedures. To support benchmarking and reproducibility, *pEYES* also provides methods for downloading and parsing publicly available, human-annotated datasets that can serve as GT. Together, these packages form an increasingly complete and collaborative open-source ecosystem, promoting transparency, reproducibility and methodological rigor in ET research.

### Study limitations

Our research evaluates the performance of seven threshold-based EM detectors. However, this analysis is limited to a single parameterization of each algorithm, using the hyper-parameters defined by Andersson et al. (2017), who published the lund2013 dataset, or those set by the creators of the *NH* (Nyström & Holmqvist, 2010) and *REMoDNaV* (Dar et al., 2021) detectors. Notably, Andersson et al. (2017) determined the saccade velocity threshold (45°/𝑠) based on the peak velocity of human-annotated saccades, whereas a more appropriate choice for the *I-VT* and *I-VVT* detectors might have been saccades’ *minimum* velocity. Adjusting velocity or dispersion thresholds alters a detector’s behavior, biasing it toward either fixation or saccade detection. Indeed, prior studies have applied different thresholds to distinguish between EM types (Birawo & Kasprowski, 2022; Blignaut, 2009; Komogortsev & Karpov, 2013; Reich et al., 2025; Salvucci & Goldberg, 2000). When applying those alternative thresholds to the *lund2013^+^-image* dataset, we observed that different threshold values led to varying proportions of detected fixations and saccades, but no single set of thresholds provided a clear advantage in terms of better aligning with the manually annotated EMs (results are available online). To maintain reproducibility, we chose to report results using the original parameterization proposed by Andersson et al., which provides decent coverage of the manually-labeled data (see **Table 3**). That said, *pEYES* allows users to revisit the results using different parameters.

Another limitation of this study is the use of human annotations as GT labels. Hessels et al. (2018) demonstrated the inherent ambiguity in the definitions and operationalizations of fixations and saccades among 120 ET researchers. Similarly, Lappi (2016) highlighted this ambiguity by reviewing how different types of EMs share overlapping physiological features and functional purposes, making them difficult to distinguish. This ambiguity introduces noise and variability into human annotations, reducing their reliability as GT labels. Hooge et al. (2018) addressed this issue directly by examining the variability in fixation classification across a dozen ET researchers. They concluded that human classification cannot be considered a true “gold standard” for fixation identification but remains a valuable tool for evaluating algorithm performance – as demonstrated in the current study.

Additionally, this study examines only a small subset of EM detection algorithms, focusing on their performance in detecting eye movements during free viewing of images. Over the past decade, the increasing adoption of machine learning (ML) and deep learning (DL) in eye-tracking research has led to the development of ML-based EM detectors (see, for example, Fuhl et al., 2021; Stratsev et al., 2019; and Zemblys et al., 2018). With the growing availability of publicly available datasets containing diverse EM types across various visual stimuli, such models have the potential to become more robust. However, existing ML-based detectors rarely outperform “classical” state-of-the-art detectors, require large-scale training data, and are often difficult to adapt for real-time classification and gaze-contingent experiments. Despite these limitations, their increasing presence in modern research suggests that future studies should incorporate ML-based detectors to enable a more comprehensive evaluation of EM detection methods.

Moving forward, *pEYES* aims to serve as a foundational tool for reproducible and transparent EM detection research. By providing standardized implementations of commonly used detectors, along with evaluation metrics and visualization tools, the package enables researchers to assess detector performance systematically and apply these insights to their own datasets. As eye-tracking research continues to evolve, integrating more advanced detection methods – including ML and DL-based approaches – will be crucial. We encourage researchers to explore, contribute to, and expand *pEYES*, which is freely available online (https://github.com/huji-hcnl/pEYES). Through continued collaboration, we hope to refine detection methodologies and further advance the field of EM research.

## Acknowledgements

We extend our gratitude to Haimasree Bhattacharya from the Center for Interdisciplinary Data Science Research (CIDR) at the Hebrew University of Jerusalem, for her assistance in publishing the *pEYES* Python package.

This study was partially supported by scholarships awarded to JN from the Israeli Smart Transportation Research Center (ISTRC) and the National Road Safety Authority of Israel, and by grants 1902/14 and 3504/20 from the Israel Science Foundation awarded to LYD. LYD is also supported by the Jack H. Skirball Research Fund. LYD is also supported by the Jack H. Skirball research fund. LYD is a founder and chief science officer of Innereye, a startup in the neurotech business which has no connection to this study.

## Open Practices Statement

The *pEYES* package is implemented in Python, a free and open-source programming language, and can be downloaded at no cost through the Python Package Index (PyPi: https://pypi.org/project/peyes/). We encourage researchers to incorporate *pEYES* into their analyses, without the need to obtain additional, often expensive, licensed software. In addition, the package’s source code, user manuals, and all code for the analyses presented in this report, are publicly available on GitHub (https://github.com/huji-hcnl/pEYES). We welcome collaboration to extend and improve the package’s functionality.

## Footnotes

1 While the term eye movement *detection* is widely used in the literature, it is technically a misnomer. As noted by Hessels et al. (2018, Section 1.4), eye-trackers do not directly measure oculomotor events but rather record features (e.g., gaze position, pupil size) from which such events are computationally inferred. Thus, algorithms do not truly *detect* events but rather impose a computational definition that *classifies* segments of the signal as fixations, saccades, etc. Nevertheless, we use the term detection throughout this article to remain consistent with prevailing usage in the field.

2 We define FAr as the ratio of *false-alarms* – i.e., unmatched detected event boundaries – to the number of *negative windows* in the GT. This definition can yield FAr values exceeding 1, particularly when the user-defined Δ𝑡 parameter is large enough to reduce the number of *negative windows* below the number of FAs. In such cases, users should apply a correction, either by truncating FAr at 1 or by applying another normalization method. An alternative approach defines *correct rejections* (*CRs*) as event boundaries of other types that were successfully matched. The FAr is then computed as 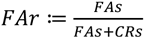. This approach avoids the risk of FAr exceeding 1, but requires detecting and matching all event types. For example, to evaluate a detector’s sensitivity to saccade onsets, the number of CRs must be derived from successfully matched fixation onsets, blink onsets, and so on. Conversely, the method used here can be applied even when only a single type of event – or even just a single event boundary (e.g., saccade onset) – is detected. Notably, both approaches are implemented in the *pEYES* package, providing users with flexibility in how detection sensitivity is evaluated.

3 Seven of the 63 recordings (42,770 samples, accounting for 11.2% of the data) were recorded at a sampling rate of 200𝐻𝑧. Of those, four recording (7985 samples) feature an image stimulus and the remaining three recordings (34,785 samples) feature a video stimulus.

4 Due to the inconsistencies in sampling rate between recordings, we measure temporal differences, *RTO*s and *RTD*s in units of samples instead of milliseconds. The results reported here were replicated when analyzing only the sixteen (out of 20) recordings that were recorded at a uniform sampling rate of 500𝐻𝑧.

5 Unlike other analyses in this report, which were performed at the level of individual recordings, temporal alignment results were aggregated over trials, yielding hundreds of measurements per detector. This may contribute to the high statistical significance observed in the Kruskal-Wallis and post-hoc tests.

