## Appendix for "Systematic Classification Differences Across Eye-Movement Detection Algorithms"

### Appendices

#### **Appendix A:** Example Visualization Implemented in *pEYES*

The *pEYES* package provides built-in visualizations for both raw ET data and parsed EM events. Below are example visualizations illustrating different aspects of the data, generated using *pEYES*.

##### **Appendix A1:** Visualizations for a Single Recording

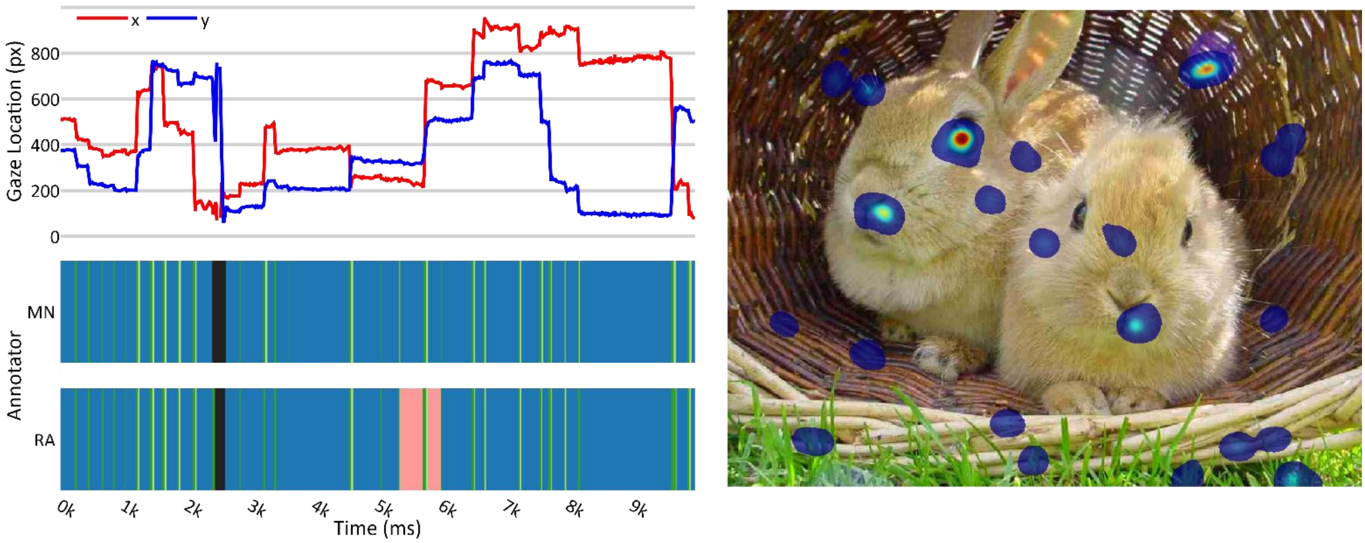


*Note:* Example visualizations from a single recording of the *lund2013⁺-image* dataset: gaze position over time (pixel coordinates; top left), scarf plots showing sample-by-sample annotations from the dataset's two human annotators, *RA* and *MN* (bottom left), and heatmap of gaze locations overlayed on the presented stimulus image (right).

**Appendix A2:** Visualizations for Multiple EM Types

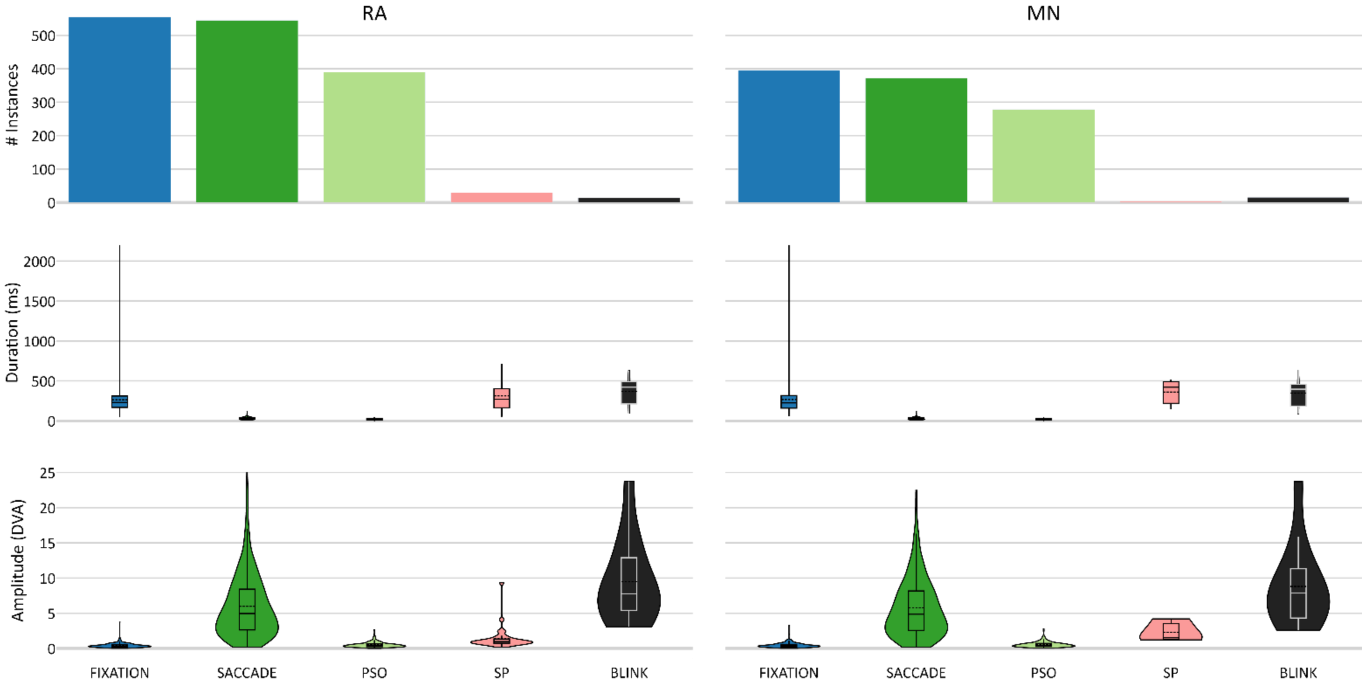


*Note:* Feature distributions across all EM events identified by human annotators *RA* and *MN* (left and right columns, respectively) in the *lund2013⁺-image* dataset, aggregated by event type (x-axis): number of detected events, event durations, and amplitudes (top, middle, and bottom rows, respectively).

**Appendix A3:** Visualization for a Single EM Type

*Note:* Main sequence plot for all saccades identified by annotator *RA* in the *lund2013⁺-image* dataset, illustrating the relationship between saccade amplitude (x-axis) and duration (y-axis). A corresponding plot for annotator *MN* is available online.

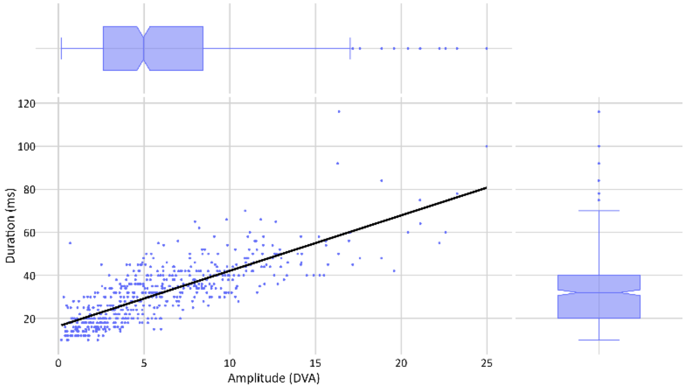

#### **Appendix B:** Detecting EMs Using *pEYES*

The pEYES package is designed for ET researchers and requires only basic Python knowledge. The code snippet below demonstrates a full EM detection pipeline: downloading a dataset, instantiating a `Detector` object, labeling raw gaze data, segmenting it into `Event` objects, and generating a basic visualization of the results.

Additional features and functionalities are described in the user manuals, available at [https://github.com/huji-hcnl/pEYES/tree/main/docs/User%20Guide](https://github.com/huji-hcnl/pEYES/tree/main/docs/User%20Guide).

```
import peyes

# load the lund2013 dataset
dataset = peyes.datasets.lund2013()

# extract single-trial data
trial1 = dataset[dataset[peyes.constants.TRIAL_ID_STR] == 1]
ps = trial1["pixel_size"].values[0]
vd = trial1["viewer_distance"].values[0]

# create a detector object
det = peyes.create_detector(
        "engbert",
        missing_value=np.nan,
        min_event_duration=4,      # in ms
        pad_blinks_time=0,         # in ms
)

# assign labels
labels, metadata = det.detect(
    t=trial1 [peyes.constants.T].values,
    x=trial1 [peyes.constants.X].values,
    y=trial1 [peyes.constants.Y].values,
    pixel_size_cm=ps,
    viewer_distance_cm=vd,
)

# generate Event objects
events = peyes.create_events(
    labels=trial1_labels,
    t=trial1_data[peyes.constants.T].values,
    x=trial1_data[peyes.constants.X].values,
    y=trial1_data[peyes.constants.Y].values,
    pupil=trial1_data[peyes.constants.PUPIL].values,
    pixel_size=trial1_pixel_size,
    viewer_distance=trial1_viewer_distance
)

# plot event features
fig = peyes.visualize.event_summary(
        events,
        show_outliers=False
)
```

### **Appendix C:** Detection Parameters

**Appendix C1:** List of Event-Specific Parameters Used in the Detection Pipeline

| Parameter Name | Value | Unit | Value From | Notes |
|---|---|---|---|---|
| min_event_samples | 2 | samples | | |
| min_event_duration | 4 | ms | | |
| pad_blinks_ms | 0 | ms | | Dar et al. (2021) used $10ms$ |
| min_fixation_duration | 55 | ms | Andersson et al. (2017) | |
| max_fixation_duration | 2500 | ms | | |
| min_saccade_duration | 10 | ms | Dar et al. (2021); Nyström & Holmqvist (2010) | |
| max_saccade_duration | 200 | ms | | |
| min_pso_duration | 4 | ms | | |
| max_pso_duration | 40 | ms | Dar et al. (2021) | |
| min_sp_duration | 40 | ms | Dar et al. (2021) | |
| max_sp_duration | 5000 | ms | | |
| min_blink_duration | 20 | ms | Dar et al. (2021) | |
| max_blink_duration | 2500 | ms | | |

**Appendix C2:** List of Algorithm-Specific Parameters Used in the Detection Pipeline

| Algorithm | Parameter Name | Value | Unit | Value From | Notes |
|---|---|---|---|---|---|
| I-VT | saccade_velocity_threshold | 45 | DVA/s | Andersson et al. (2017) | Hooge et al. (2018) used $16.5\ {}^{\circ}/_{s}$ |
| I-VVT | saccade_velocity_threshold | 45 | DVA/s | Andersson et al. (2017) | |
| | smooth_pursuit_velocity_threshold | 26 | DVA/s | Komogortsev & Karpov (2013) | |
| I-DT | dispersion_threshold | 2.7 | DVA | Andersson et al. (2017) | Salvucci & Goldberg (2000) used $0.5\text{-}1°$ |
| | window_duration | 55 | ms | Andersson et al. (2017) | Salvucci & Goldberg (2000) used $100ms$ |
| I-DVT | saccade_velocity_threshold | 45 | DVA/s | Andersson et al. (2017) | |
| | dispersion_threshold | 2.7 | DVA | Andersson et al. (2017) | Salvucci & Goldberg (2000) used $0.5\text{-}1°$ |
| | window_duration | 55 | ms | Andersson et al. (2017) | Salvucci & Goldberg (2000) used $100ms$ |
| Engbert | lambda | 6 | | Andersson et al. (2017); Engbert & Mergenthaler (2006) | Engbert & Kliegl (2003) used 5 |
| | derivation_window_size | 5 | samples | Andersson et al. (2017); Engbert & Mergenthaler (2006) | |

| Algorithm | Parameter Name | Value | Unit | Value From | Notes |
|---|---|---|---|---|---|
| NH | filter_duration_ms | 20 | ms | Nyström & Holmqvist (2010) | $2 \times$ min_saccade_duration |
| | filter_polyorder | 2 | | Nyström & Holmqvist (2010) | |
| | saccade_max_velocity | 1000 | DVA/s | Nyström & Holmqvist (2010) | |
| | saccade_max_acceleration | 100000 | DVA/$s^2$ | Nyström & Holmqvist (2010) | |
| | alpha | 0.7 | | Nyström & Holmqvist (2010) | |
| | beta | 0.3 | | Nyström & Holmqvist (2010) | |
| | median_filter_duration_ms | 50 | ms | Dar et al. (2021) | argument name in original paper: median_filter_length |
| | savgol_filter_polyorder | 2 | | Dar et al. (2021) | argument name in original paper: savgol_polyord |
| | median_filter_duration_ms | 50 | ms | Dar et al. (2021) | argument name in original paper: median_filter_length |
| | savgol_filter_polyorder | 2 | | Dar et al. (2021) | argument name in original paper: savgol_polyord |
| | savgol_filter_duration_ms | 19 | ms | Dar et al. (2021) | argument name in original paper: savgol_length |
| | max_velocity | 1500 | DVA/s | | argument name in original paper: max_vel<br>argument value in original paper: $1000 {}^{\circ}/_{s}$ |
| | min_intersaccade_duration | 20 | ms | | minimum fixation/sp/blink duration<br>argument value in original paper: $40ms$ |

| Algorithm | Parameter Name | Value | Unit | Value From | Notes |
|---|---|---|---|---|---|
| REMoDNaV | saccade_onset_threshold_noise_factor | 5 | | Dar et al. (2021) | argument name in original paper: noise_factor |
| | saccade_initial_velocity_threshold | 300 | DVA/s | Dar et al. (2021); Nyström & Holmqvist (2010) | argument name in original paper: velthresh_startvelocity |
| | saccade_initial_max_freq | 2 | Hz | Dar et al. (2021) | argument name in original paper: max_initial_saccade_freq |
| | saccade_context_window_duration | 1000 | ms | Dar et al. (2021) | argument name in original paper: saccade_context_window_length |
| | smooth_pursuits_lowpass_cutoff_freq | 4 | Hz | Dar et al. (2021) | argument name in original paper: lowpass_cutoff_freq |
| | smooth_pursuit_drift_velocity_threshold | 2 | DVA/s | Dar et al. (2021) | argument name in original paper: pursuit_velthresh |

#### **Appendix D:** Sample-by-Sample Agreement Scores

We statistically compared detector agreement with each human annotator separately, using *Cohen's Kappa*, *MCC*, and *1-NLD* as agreement measures. For each annotator, we first applied a Friedman test to assess overall differences across detectors, followed by post-hoc pairwise comparisons using the Tukey-HSD test, which accounts for multiple comparisons. The resources below provide detailed results, specifying which human annotator was used as the GT for each analysis. For pairwise comparison tables, the top section indicates significance levels, and the bottom section shows the corrected p-values. Significance is denoted as follows:

$$\dagger\!: p < 0.075, \quad *\!: p < 0.05, \quad **\!: p < 0.01, \quad ***\!: p < 0.001, \quad n.s.: \text{not significant}$$

**Appendix D1:** Distribution of Agreement Scores Between Annotator *MN* and Detectors

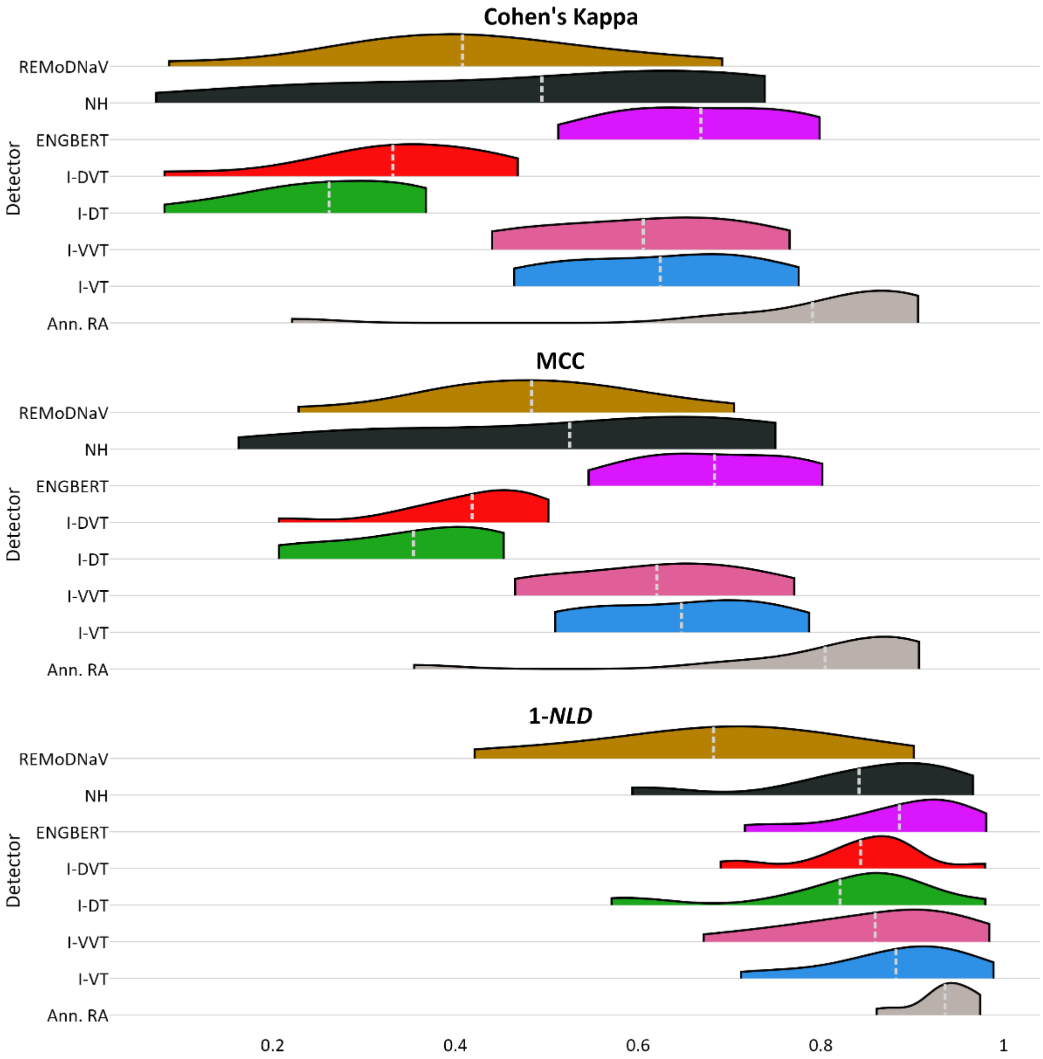


*Note:* Similar to **Figure 2**, this figure shows the distribution of sample-by-sample agreement scores (*Cohen's Kappa*, *MCC*, *1-NLD*) between GT annotator *MN* and each detector, across recordings. Inter-rater agreement (with annotator *RA*) is provided for comparison.

**Appendix D2:** Friedman Test Results

| | ***Cohen's Kappa*** | | ***MCC*** | | ***1-NLD*** | |
|---|---|---|---|---|---|---|
| | $Q(6)$ | $p$ | $Q(6)$ | $p$ | $Q(6)$ | $p$ |
| ***RA*** | 76.5 | $< 0.001$ | 81.8 | $< 0.001$ | 80.2 | $< 0.001$ |
| ***MN*** | 61.6 | $< 0.001$ | 58.6 | $< 0.001$ | 62.8 | $< 0.001$ |

*Note:* Results of Friedman tests comparing agreement scores across the seven detectors ($df = 6$), conducted separately using each human annotator (*RA* and *MN*) as ground truth.

**Appendix D3:** Pairwise Comparison Results (*Cohen's Kappa* Scores)

| | | ivt | ivvt | idt | idvt | engbert | nh | remodnav |
|---|---|---|---|---|---|---|---|---|
| ivt | MN | -- | n.s. | *** | * | n.s. | n.s. | n.s. |
| | RA | -- | n.s. | *** | ** | n.s. | n.s. | n.s. |
| ivvt | MN | 1.0000 | -- | ** | * | n.s. | n.s. | n.s. |
| | RA | 1.0000 | -- | *** | ** | n.s. | n.s. | n.s. |
| idt | MN | 0.0006 | 0.0013 | -- | n.s. | *** | n.s. | n.s. |
| | RA | 0.0003 | 0.0003 | -- | n.s. | *** | † | n.s. |
| idvt | MN | 0.0140 | 0.0259 | 0.9923 | -- | ** | n.s. | n.s. |
| | RA | 0.0052 | 0.0048 | 0.9969 | -- | *** | n.s. | n.s. |
| engbert | MN | 0.9973 | 0.9894 | 0.0000 | 0.0010 | -- | n.s. | * |
| | RA | 0.9855 | 0.9872 | 0.0000 | 0.0001 | -- | n.s. | n.s. |
| nh | MN | 0.8163 | 0.8926 | 0.1225 | 0.5188 | 0.4215 | -- | n.s. |
| | RA | 0.8914 | 0.8839 | 0.0519 | 0.2580 | 0.3880 | -- | n.s. |
| remodnav | MN | 0.1859 | 0.2683 | 0.7186 | 0.9832 | 0.0331 | 0.9553 | -- |
| | RA | 0.7795 | 0.7685 | 0.1030 | 0.3974 | 0.2505 | 1.0000 | -- |

**Appendix D4:** Pairwise Comparison Results (*MCC* Scores)

| | | ivt | ivvt | idt | idvt | engbert | nh | remodnav |
|---|---|---|---|---|---|---|---|---|
| ivt | MN | -- | n.s. | *** | * | n.s. | n.s. | n.s. |
| | RA | -- | n.s. | *** | ** | n.s. | n.s. | n.s. |
| ivvt | MN | 0.9998 | -- | ** | † | n.s. | n.s. | n.s. |
| | RA | 1.0000 | -- | ** | * | n.s. | n.s. | n.s. |
| idt | MN | 0.0005 | 0.0031 | -- | n.s. | *** | n.s. | n.s. |
| | RA | 0.0003 | 0.0012 | -- | n.s. | *** | n.s. | n.s. |
| idvt | MN | 0.0152 | 0.0562 | 0.9898 | -- | ** | n.s. | n.s. |
| | RA | 0.0094 | 0.0235 | 0.9927 | -- | *** | n.s. | n.s. |
| engbert | MN | 0.9985 | 0.9729 | 0.0000 | 0.0015 | -- | n.s. | † |
| | RA | 0.9923 | 0.9656 | 0.0000 | 0.0003 | -- | n.s. | n.s. |
| nh | MN | 0.7117 | 0.9047 | 0.1836 | 0.6582 | 0.3442 | -- | n.s. |
| | RA | 0.6997 | 0.8385 | 0.1596 | 0.5864 | 0.2309 | -- | n.s. |
| remodnav | MN | 0.2342 | 0.4659 | 0.6398 | 0.9729 | 0.0562 | 0.9916 | -- |
| | RA | <0.0001 | <0.0001 | 0.8658 | 0.4016 | <0.0001 | 0.0020 | -- |

**Appendix D5:** Pairwise Comparison Results (*1-NLD* Scores)

| | | ivt | ivvt | idt | idvt | engbert | nh | remodnav |
|---|---|---|---|---|---|---|---|---|
| ivt | MN | -- | n.s. | n.s. | n.s. | n.s. | n.s. | * |

| | | | | | | | | |
|---|---|---|---|---|---|---|---|---|
| | RA | -- | n.s. | n.s. | n.s. | n.s. | n.s. | ** |
| ivvt | MN | 0.9985 | -- | n.s. | n.s. | n.s. | n.s. | † |
| | RA | 0.9992 | -- | n.s. | n.s. | n.s. | n.s. | * |
| idt | MN | 0.8511 | 0.9879 | -- | n.s. | n.s. | n.s. | n.s. |
| | RA | 0.5094 | 0.8235 | -- | n.s. | n.s. | n.s. | n.s. |
| idvt | MN | 0.9236 | 0.9973 | 1.0000 | -- | n.s. | n.s. | n.s. |
| | RA | 0.6152 | 0.8914 | 1.0000 | -- | n.s. | n.s. | n.s. |
| engbert | MN | 1.0000 | 0.9932 | 0.7602 | 0.8586 | -- | n.s. | ** |
| | RA | 1.0000 | 0.9931 | 0.3594 | 0.4596 | -- | n.s. | ** |
| nh | MN | 0.9904 | 1.0000 | 0.9979 | 0.9998 | 0.9721 | -- | n.s. |
| | RA | 0.9543 | 0.9986 | 0.9823 | 0.9938 | 0.8864 | -- | n.s. |
| remodnav | MN | 0.0113 | 0.0726 | 0.4325 | 0.3130 | 0.0057 | 0.1324 | -- |
| | RA | 0.0035 | 0.0243 | 0.6141 | 0.5083 | 0.0013 | 0.1269 | -- |

#### **Appendix E:** Sample-by-Sample Sensitivity Scores ($d'$)

We statistically compared detector sample-level sensitivity indices ($d'$) for fixations and saccades, treating each human annotator (*RA* and *MN*) as GT in separate analyses. For each annotator, we first used a Friedman test to assess overall differences across detectors. This was followed by post-hoc pairwise comparisons using the Tukey-HSD test, which accounts for multiple comparisons. The results provided below indicate which annotator was used as GT in each analysis. For pairwise comparison tables, the top section indicates significance levels, and the bottom section shows the corrected p-values. Significance is denoted as follows:

$$\dagger: p < 0.075, \quad *: p < 0.05, \quad **: p < 0.01, \quad ***: p < 0.001, \quad n.s.: \text{not significant}$$

##### **Appendix E1:** Distribution of Sample-Level Sensitivity Scores ($d'$)

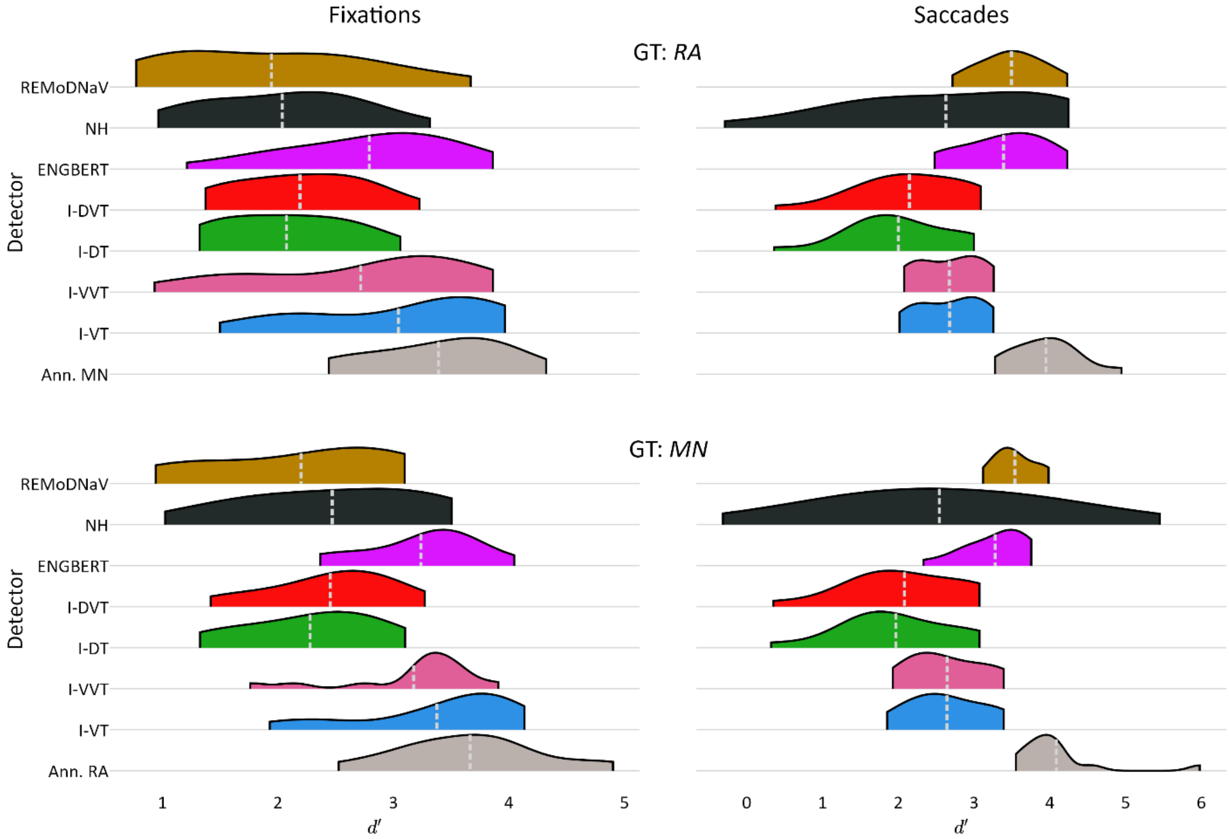


*Note:* Distribution of sample-level sensitivity indices ($d'$) across recordings for fixation (left column) and saccade (right column) detection. Each row corresponds to a different human annotator used as ground truth: *RA* (top) and *MN* (bottom). For each case, sensitivity scores of all detectors are shown, including the 2nd annotator for reference. The dashed light-gray line denotes the distribution mean.

**Appendix E2:** Friedman Test Results

| | **Fixation** | | **Saccade** | |
|---|---|---|---|---|
| | $Q(6)$ | $p$ | $Q(6)$ | $p$ |
| ***RA*** | $70.5$ | $< 0.001$ | $78.6$ | $< 0.001$ |
| ***MN*** | $54.0$ | $< 0.001$ | $56.2$ | $< 0.001$ |

*Note:* Results of Friedman tests comparing sensitivity indices ($d'$) across the seven detectors ($df = 6$), conducted separately using each human annotator (*RA* and *MN*) as ground truth.

**Appendix E3:** Pairwise Comparison Results of Sample-Level **Fixation** Sensitivity Scores ($d'$)

| | | ivt | ivvt | idt | idvt | engbert | nh | remodnav |
|---|---|---|---|---|---|---|---|---|
| ivt | MN | -- | n.s. | * | † | n.s. | n.s. | * |
| | RA | -- | n.s. | * | n.s. | n.s. | * | * |
| ivvt | MN | 0.9982 | -- | n.s. | n.s. | n.s. | n.s. | n.s. |
| | RA | 0.9751 | -- | n.s. | n.s. | n.s. | n.s. | n.s. |
| idt | MN | 0.0144 | 0.0918 | -- | n.s. | † | n.s. | n.s. |
| | RA | 0.0383 | 0.3756 | -- | n.s. | n.s. | n.s. | n.s. |
| idvt | MN | 0.0669 | 0.2727 | 0.9995 | -- | n.s. | n.s. | n.s. |
| | RA | 0.1333 | 0.6661 | 0.9996 | -- | n.s. | n.s. | n.s. |
| engbert | MN | 0.9997 | 1.0000 | 0.0606 | 0.2016 | -- | n.s. | † |
| | RA | 0.9950 | 1.0000 | 0.2389 | 0.5027 | -- | n.s. | n.s. |
| nh | MN | 0.1390 | 0.4362 | 0.9930 | 1.0000 | 0.3442 | -- | n.s. |
| | RA | 0.0286 | 0.3221 | 1.0000 | 0.9989 | 0.1977 | -- | n.s. |
| remodnav | MN | 0.0137 | 0.0884 | 1.0000 | 0.9994 | 0.0582 | 0.9923 | -- |
| | RA | 0.0120 | 0.1977 | 0.9999 | 0.9912 | 0.1099 | 1.0000 | -- |

**Appendix E4:** Pairwise Comparison Results of Sample-Level **Saccade** Sensitivity Scores ($d'$)

| | | ivt | ivvt | idt | idvt | engbert | nh | remodnav |
|---|---|---|---|---|---|---|---|---|
| ivt | MN | -- | n.s. | n.s. | n.s. | n.s. | n.s. | n.s. |
| | RA | -- | n.s. | n.s. | n.s. | n.s. | n.s. | † |
| ivvt | MN | 1.0000 | -- | n.s. | n.s. | n.s. | n.s. | n.s. |
| | RA | 1.0000 | -- | n.s. | n.s. | n.s. | n.s. | † |
| idt | MN | 0.7356 | 0.7422 | -- | n.s. | ** | n.s. | *** |
| | RA | 0.4640 | 0.4728 | -- | n.s. | *** | n.s. | *** |
| idvt | MN | 0.8729 | 0.8774 | 1.0000 | -- | * | n.s. | *** |
| | RA | 0.7657 | 0.7731 | 0.9995 | -- | *** | n.s. | *** |
| engbert | MN | 0.5036 | 0.4960 | 0.0073 | 0.0201 | -- | n.s. | n.s. |
| | RA | 0.1717 | 0.1665 | 0.0001 | 0.0007 | -- | n.s. | n.s. |
| nh | MN | 1.0000 | 1.0000 | 0.7322 | 0.8706 | 0.5074 | -- | n.s. |
| | RA | 0.9999 | 0.9999 | 0.2513 | 0.5373 | 0.3514 | -- | n.s. |
| remodnav | MN | 0.0925 | 0.0898 | 0.0002 | 0.0006 | 0.9863 | 0.0940 | -- |
| | RA | 0.0623 | 0.0635 | 0.9998 | 0.9624 | 0 | 0.0453 | -- |

#### **Appendix F:** Fixation & Saccade Temporal Alignment

We statistically compared detectors' temporal misalignment with each human annotator (*RA* and *MN*). For each annotator, we used a Kruskal-Wallis test to assess overall differences across detectors, followed by post-hoc pairwise comparisons using Dunn's test with Bonferroni correction for multiple comparisons. Importantly, the analyses depicted here were performed across all detected fixations or saccades, yielding hundreds of measurements per detector. These large sample sizes may have contributed to the high significance observed in subsequent statistical tests.
The results provided below indicate which annotator was used as GT in each analysis. For pairwise comparison tables, the top section indicates significance levels, and the bottom section shows the corrected p-values. Significance is denoted as follows:

$$\dagger: p < 0.075, \quad *: p < 0.05, \quad **: p < 0.01, \quad ***: p < 0.001, \quad n.s.: \text{not significant}$$

##### **Appendix F1:** Distribution of Temporal Misalignments Relative to Fixation and Saccade Boundaries Identified by MN

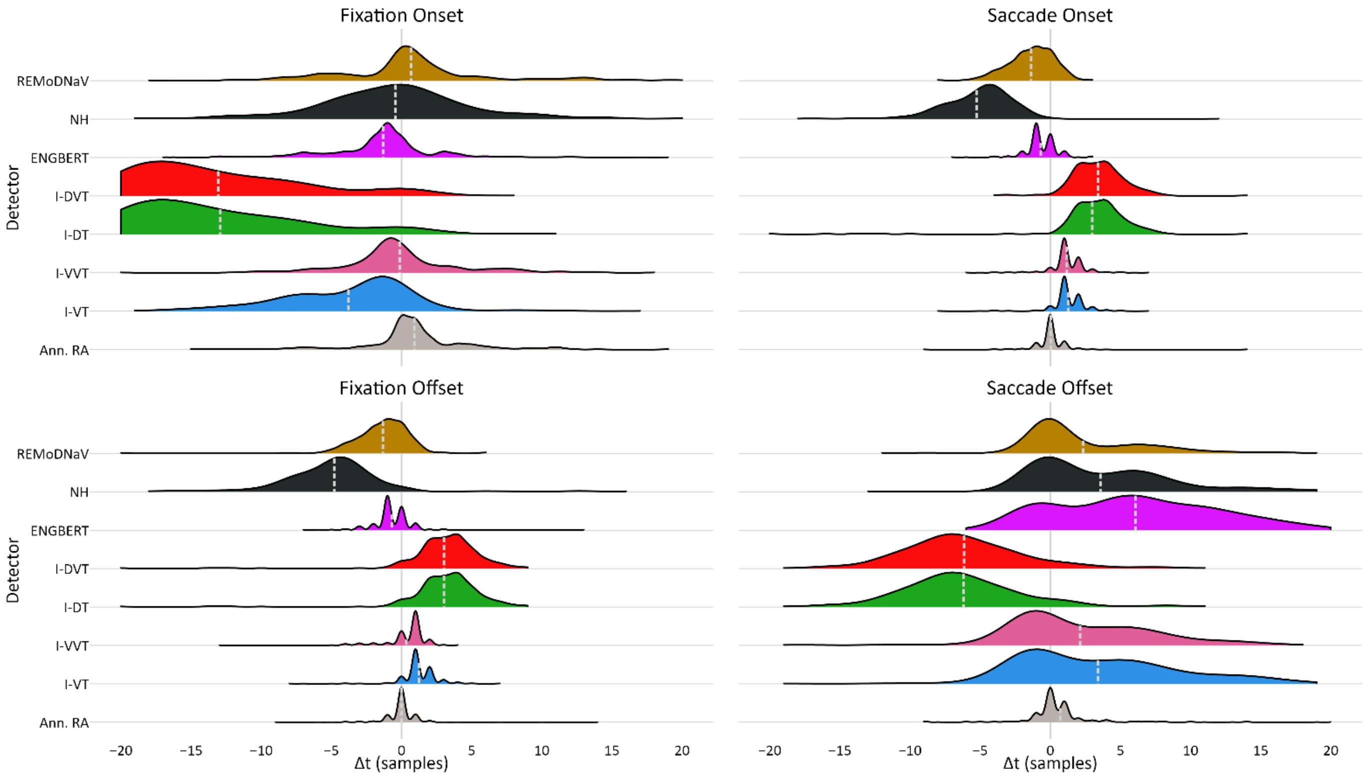


*Note:* Similar to **Figure 3**, this figure shows the distribution of temporal misalignments (in samples) between detected fixation and saccade onsets and offsets that were matched to corresponding events annotated by *MN*. Misalignment values were truncated to the range $[-20, 20]$ $samples$. The mean of each distribution, marked by a light gray dashed line, is the detector's *RTO*, and the standard deviation reflects its *RTD*. Both values are specified in **Table 4**.

**Appendix F2:** Kruskal-Wallis Test Results

| | **Fixation Onset** | | **Fixation Offset** | | **Saccade Onset** | | **Saccade Offset** | |
|---|---|---|---|---|---|---|---|---|
| | $H(6)$ | $p$ | $H(6)$ | $p$ | $H(6)$ | $p$ | $H(6)$ | $p$ |
| ***RA*** | $997.7$ | $< 0.001$ | $2070.3$ | $< 0.001$ | $2375.9$ | $< 0.001$ | $1301.5$ | $< 0.001$ |
| ***MN*** | $751.9$ | $< 0.001$ | $1394.4$ | $< 0.001$ | $1527.6$ | $< 0.001$ | $793.3$ | $< 0.001$ |

*Note:* Results of Kruskal-Wallis tests comparing temporal differences of paired fixation and saccade onsets and offsets, across the seven detectors ($df = 6$). Tests were conducted separately for each human annotator (*RA* and *MN*) as ground truth.

**Appendix F3:** Pairwise Comparison Results of **Fixation Onset** Temporal Alignments

| | | ivt | ivvt | idt | idvt | engbert | nh | remodnav |
|---|---|---|---|---|---|---|---|---|
| ivt | MN | -- | *** | *** | *** | *** | *** | *** |
| | RA | -- | *** | *** | *** | *** | *** | *** |
| ivvt | MN | <0.0001 | -- | *** | *** | * | n.s. | n.s. |
| | RA | <0.0001 | -- | *** | *** | * | n.s. | * |
| idt | MN | <0.0001 | <0.0001 | -- | n.s. | *** | *** | *** |
| | RA | <0.0001 | <0.0001 | -- | n.s. | *** | *** | *** |
| idvt | MN | <0.0001 | <0.0001 | 1.0000 | -- | *** | *** | *** |
| | RA | <0.0001 | <0.0001 | 1.0000 | -- | *** | *** | *** |
| engbert | MN | <0.0001 | 0.0145 | <0.0001 | <0.0001 | -- | n.s. | *** |
| | RA | <0.0001 | 0.0197 | <0.0001 | <0.0001 | -- | ** | *** |
| nh | MN | <0.0001 | 1.0000 | <0.0001 | <0.0001 | 0.3212 | -- | † |
| | RA | <0.0001 | 1.0000 | <0.0001 | <0.0001 | 0.0064 | -- | n.s. |
| remodnav | MN | <0.0001 | 0.1752 | <0.0001 | <0.0001 | <0.0001 | 0.0618 | -- |
| | RA | <0.0001 | 0.0144 | <0.0001 | <0.0001 | <0.0001 | 0.2296 | -- |

**Appendix F4:** Pairwise Comparison Results of **Fixation Offset** Temporal Alignments

| | | ivt | ivvt | idt | idvt | engbert | nh | remodnav |
|---|---|---|---|---|---|---|---|---|
| ivt | MN | -- | *** | *** | *** | *** | *** | *** |
| | RA | -- | *** | *** | *** | *** | *** | *** |
| ivvt | MN | <0.0001 | -- | *** | *** | *** | *** | *** |
| | RA | <0.0001 | -- | *** | *** | *** | *** | *** |
| idt | MN | <0.0001 | <0.0001 | -- | n.s. | *** | *** | *** |
| | RA | <0.0001 | <0.0001 | -- | n.s. | *** | *** | *** |
| idvt | MN | <0.0001 | <0.0001 | 1.0000 | -- | *** | *** | *** |
| | RA | <0.0001 | <0.0001 | 1.0000 | -- | *** | *** | *** |
| engbert | MN | <0.0001 | <0.0001 | <0.0001 | <0.0001 | -- | *** | n.s. |
| | RA | <0.0001 | <0.0001 | <0.0001 | <0.0001 | -- | *** | n.s. |
| nh | MN | <0.0001 | <0.0001 | <0.0001 | <0.0001 | <0.0001 | -- | *** |
| | RA | <0.0001 | <0.0001 | <0.0001 | <0.0001 | <0.0001 | -- | *** |

| | | | | | | | | |
|---|---|---|---|---|---|---|---|---|
| remodnav | MN | <0.0001 | <0.0001 | <0.0001 | <0.0001 | 0.9408 | <0.0001 | -- |
| | RA | <0.0001 | <0.0001 | <0.0001 | <0.0001 | 0.1623 | <0.0001 | -- |

**Appendix F5:** Pairwise Comparison Results of **Saccade Onset** Temporal Alignments

| | | ivt | ivvt | idt | idvt | engbert | nh | remodnav |
|---|---|---|---|---|---|---|---|---|
| ivt | MN | -- | n.s. | *** | *** | *** | *** | *** |
| | RA | -- | n.s. | *** | *** | *** | *** | *** |
| ivvt | MN | 1.0000 | -- | *** | *** | *** | *** | *** |
| | RA | 1.0000 | -- | *** | *** | *** | *** | *** |
| idt | MN | <0.0001 | <0.0001 | -- | n.s. | *** | *** | *** |
| | RA | <0.0001 | <0.0001 | -- | n.s. | *** | *** | *** |
| idvt | MN | <0.0001 | <0.0001 | 1.0000 | -- | *** | *** | *** |
| | RA | <0.0001 | <0.0001 | 1.0000 | -- | *** | *** | *** |
| engbert | MN | <0.0001 | <0.0001 | <0.0001 | <0.0001 | -- | *** | n.s. |
| | RA | <0.0001 | <0.0001 | <0.0001 | <0.0001 | -- | *** | n.s. |
| nh | MN | <0.0001 | <0.0001 | <0.0001 | <0.0001 | <0.0001 | -- | *** |
| | RA | <0.0001 | <0.0001 | <0.0001 | <0.0001 | <0.0001 | -- | *** |
| remodnav | MN | <0.0001 | <0.0001 | <0.0001 | <0.0001 | 0.1512 | <0.0001 | -- |
| | RA | <0.0001 | <0.0001 | <0.0001 | <0.0001 | 0.0903 | <0.0001 | -- |

**Appendix F6:** Pairwise Comparison Results of **Saccade Offset** Temporal Alignments

| | | ivt | ivvt | idt | idvt | engbert | nh | remodnav |
|---|---|---|---|---|---|---|---|---|
| ivt | MN | -- | n.s. | *** | *** | *** | n.s. | n.s. |
| | RA | -- | n.s. | *** | *** | *** | n.s. | n.s. |
| ivvt | MN | 0.3596 | -- | *** | *** | *** | * | n.s. |
| | RA | 0.4770 | -- | *** | *** | *** | ** | n.s. |
| idt | MN | <0.0001 | <0.0001 | -- | n.s. | *** | *** | *** |
| | RA | <0.0001 | <0.0001 | -- | n.s. | *** | *** | *** |
| idvt | MN | <0.0001 | <0.0001 | 1.0000 | -- | *** | *** | *** |
| | RA | <0.0001 | <0.0001 | 1.0000 | -- | *** | *** | *** |
| engbert | MN | <0.0001 | <0.0001 | <0.0001 | <0.0001 | -- | ** | *** |
| | RA | <0.0001 | <0.0001 | <0.0001 | <0.0001 | -- | *** | *** |
| nh | MN | 1.0000 | 0.0181 | <0.0001 | <0.0001 | 0.0016 | -- | n.s. |
| | RA | 1.0000 | 0.0013 | <0.0001 | <0.0001 | 0.0001 | -- | n.s. |
| remodnav | MN | 1.0000 | 1.0000 | <0.0001 | <0.0001 | <0.0001 | 0.7445 | -- |
| | RA | 1.0000 | 1.0000 | <0.0001 | <0.0001 | <0.0001 | 0.2968 | -- |

#### **Appendix G:** Fixation Boundary Sensitivity

We statistically compared detectors' sensitivity index scores ($d'$) for fixation onsets and offsets, using each human annotator (*RA* and *MN*) as GT. For each annotator, we first applied a Friedman test to assess overall differences across detectors, followed by post-hoc pairwise comparisons using the Tukey-HSD test, which accounts for multiple comparisons. The results provided below indicate which annotator was used as GT in each analysis. For pairwise comparison tables, the top section indicates significance levels, and the bottom section shows the corrected p-values. Significance is denoted as follows:

$$\dagger: p < 0.075, \quad *: p < 0.05, \quad **: p < 0.01, \quad ***: p < 0.001, \quad n.s.: \text{not significant}$$

##### **Appendix G1:** Fixation Boundary Sensitivity Index (d') Across Temporal Thresholds Relative to Annotator MN

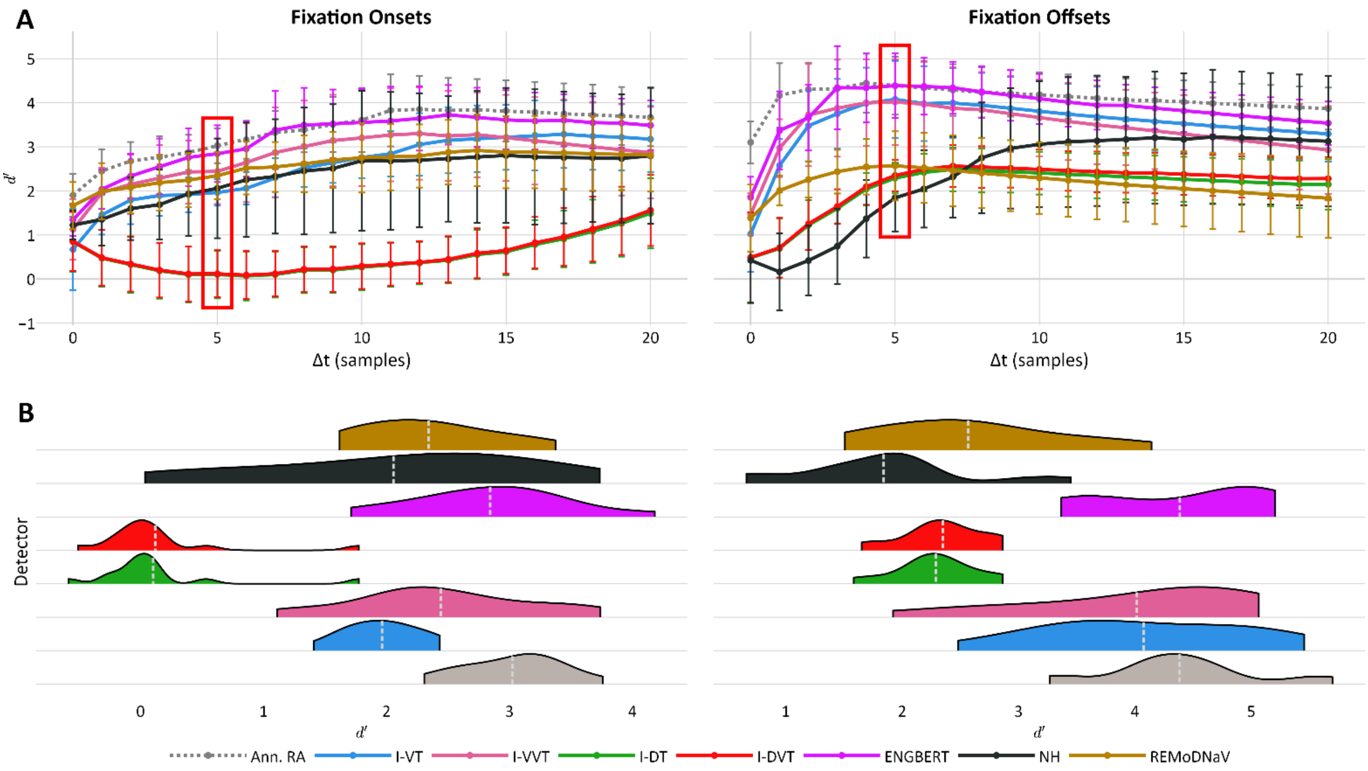


*Note:* Similar to **Figure 4**, this figure shows sensitivity index ($d'$) scores for fixation onset and offset detection by each detector, using human annotator *MN* as GT. Sensitivity scores of the second human annotator (*RA*) are shown for reference (gray line & violin).

**Appendix G2:** Friedman Test Results

| | Fixation Onset | | Fixation Offset | |
|---|---|---|---|---|
| | $Q(6)$ | $p$ | $Q(6)$ | $p$ |
| ***RA*** | 84.6 | $< 0.001$ | 79.4 | $< 0.001$ |
| ***MN*** | 66.0 | $< 0.001$ | 59.1 | $< 0.001$ |

*Note:* Results of Friedman tests comparing fixation onset and offset sensitivity indices ($d'$) across the seven detectors ($df = 6$), conducted separately using each human annotator (*RA* and *MN*) as ground truth.

**Appendix G3:** Pairwise Comparison Results of **Fixation Onset** Sensitivity Scores

| | | ivt | ivvt | idt | idvt | engbert | nh | remodnav |
|---|---|---|---|---|---|---|---|---|
| ivt | MN | -- | n.s. | n.s. | n.s. | n.s. | n.s. | n.s. |
| | RA | -- | n.s. | ** | ** | n.s. | n.s. | n.s. |
| ivvt | MN | 0.8729 | -- | ** | ** | n.s. | n.s. | n.s. |
| | RA | 0.9447 | -- | *** | *** | n.s. | n.s. | n.s. |
| idt | MN | 0.1324 | 0.0012 | -- | n.s. | *** | * | ** |
| | RA | 0.0056 | 0.0000 | -- | n.s. | *** | ** | *** |
| idvt | MN | 0.1539 | 0.0015 | 1.0000 | -- | *** | * | ** |
| | RA | 0.0066 | <0.0001 | 1.0000 | -- | *** | ** | *** |
| engbert | MN | 0.3409 | 0.9826 | <0.0001 | <0.0001 | -- | n.s. | n.s. |
| | RA | 0.4292 | 0.9726 | <0.0001 | <0.0001 | -- | n.s. | n.s. |
| nh | MN | 0.9923 | 0.9983 | 0.0127 | 0.0160 | 0.8163 | -- | n.s. |
| | RA | 0.9998 | 0.9936 | 0.0011 | 0.0013 | 0.6809 | -- | n.s. |
| remodnav | MN | 0.9598 | 1.0000 | 0.0042 | 0.0054 | 0.9277 | 1.0000 | -- |
| | RA | 0.9979 | 0.9991 | 0.0004 | 0.0005 | 0.8085 | 1.0000 | -- |

**Appendix G4:** Pairwise Comparison Results of **Fixation Offset** Sensitivity Scores

| | | ivt | ivvt | idt | idvt | engbert | nh | remodnav |
|---|---|---|---|---|---|---|---|---|
| ivt | MN | -- | n.s. | * | * | n.s. | *** | n.s. |
| | RA | -- | n.s. | ** | ** | n.s. | *** | * |
| ivvt | MN | 1.0000 | -- | * | † | n.s. | *** | n.s. |
| | RA | 0.9997 | -- | * | * | n.s. | *** | n.s. |
| idt | MN | 0.0165 | 0.0401 | -- | n.s. | ** | n.s. | n.s. |
| | RA | 0.0030 | 0.0157 | -- | n.s. | *** | n.s. | n.s. |
| idvt | MN | 0.0313 | 0.0703 | 1.0000 | -- | ** | n.s. | n.s. |
| | RA | 0.0046 | 0.0228 | 1.0000 | -- | *** | n.s. | n.s. |
| engbert | MN | 0.9996 | 0.9948 | 0.0029 | 0.0063 | -- | *** | * |
| | RA | 0.9806 | 0.8774 | <0.0001 | 0.0001 | -- | *** | *** |
| nh | MN | 0.0001 | 0.0005 | 0.9536 | 0.9017 | <0.0001 | -- | n.s. |
| | RA | <0.0001 | 0.0002 | 0.9612 | 0.9377 | <0.0001 | -- | n.s. |
| remodnav | MN | 0.0961 | 0.1836 | 0.9985 | 0.9999 | 0.0252 | 0.7186 | -- |
| | RA | 0.0299 | 0.1048 | 0.9977 | 0.9992 | 0.0008 | 0.7140 | -- |

#### **Appendix H:** Saccade Boundary Sensitivity

We statistically compared detectors' sensitivity index scores ($d'$) for saccade onsets and offsets, using each human annotator (*RA* and *MN*) as GT. For each annotator, we first applied a Friedman test to assess overall differences across detectors, followed by post-hoc pairwise comparisons using the Tukey-HSD test, which accounts for multiple comparisons. The results provided below indicate which annotator was used as GT in each analysis. For pairwise comparison tables, the top section indicates significance levels, and the bottom section shows the corrected p-values. Significance is denoted as follows:

$\dagger: p < 0.075, \quad *: p < 0.05, \quad **: p < 0.01, \quad ***: p < 0.001, \quad n.s.:$ not significant

**Appendix H1:** Saccade Boundary Sensitivity Index (d') Across Temporal Thresholds Relative to Annotator MN

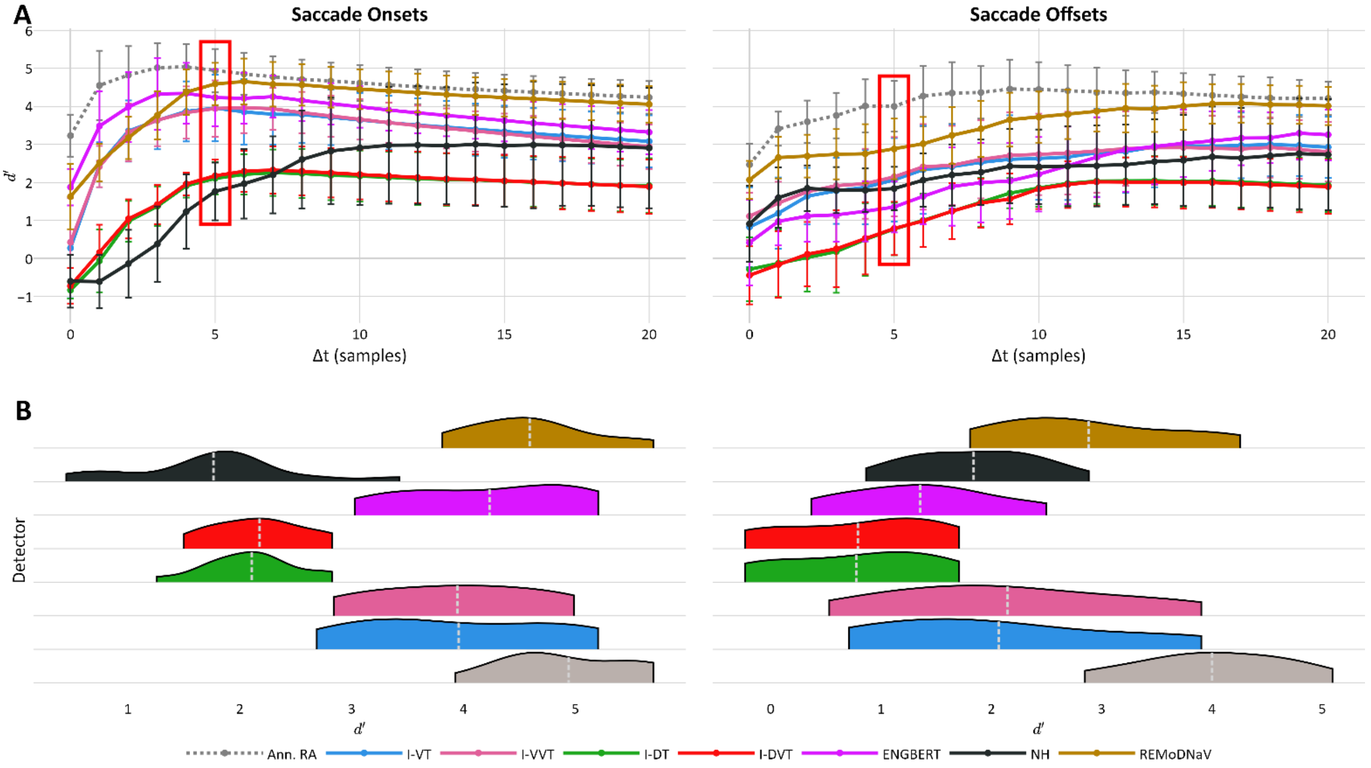


*Note: Note:* Similar to **Figure 5**, this figure shows sensitivity index ($d'$) scores for saccade onset and offset detection by each detector, using human annotator *MN* as GT. Sensitivity scores of the second human annotator (*RA*) are shown for reference (gray line & violin).

**Appendix H2:** Friedman Test Results

| | Saccade Onset | | Saccade Offset | |
|---|---|---|---|---|
| | $Q(6)$ | $p$ | $Q(6)$ | $p$ |
| ***RA*** | $86.9$ | $< 0.001$ | $73.7$ | $< 0.001$ |
| ***MN*** | $64.1$ | $< 0.001$ | $32.0$ | $< 0.001$ |

*Note:* Results of Friedman tests comparing saccade onset and offset sensitivity indices ($d'$) across the seven detectors ($df = 6$), conducted separately using each human annotator (*RA* and *MN*) as ground truth.

**Appendix H3:** Pairwise Comparison Results of **Saccade Onset** Sensitivity Scores

| | | ivt | ivvt | idt | idvt | engbert | nh | remodnav |
|---|---|---|---|---|---|---|---|---|
| ivt | MN | -- | n.s. | * | * | n.s. | ** | n.s. |
| | RA | -- | n.s. | ** | ** | n.s. | *** | n.s. |
| ivvt | MN | 1.0000 | -- | * | * | n.s. | ** | n.s. |
| | RA | 0.9999 | -- | ** | * | n.s. | *** | n.s. |
| idt | MN | 0.0179 | 0.0205 | -- | n.s. | ** | n.s. | *** |
| | RA | 0.0010 | 0.0049 | -- | n.s. | *** | n.s. | *** |
| idvt | MN | 0.0285 | 0.0325 | 1.0000 | -- | ** | n.s. | *** |
| | RA | 0.0025 | 0.0108 | 1.0000 | -- | *** | n.s. | *** |
| engbert | MN | 0.9994 | 0.9991 | 0.0027 | 0.0048 | -- | *** | n.s. |
| | RA | 0.9952 | 0.9567 | <0.0001 | 0.0001 | -- | *** | n.s. |
| nh | MN | 0.0020 | 0.0024 | 0.9987 | 0.9956 | 0.0002 | -- | *** |
| | RA | 0.0001 | 0.0006 | 0.9996 | 0.9965 | <0.0001 | -- | *** |
| remodnav | MN | 0.9704 | 0.9639 | 0.0003 | 0.0005 | 0.9993 | <0.0001 | -- |
| | RA | 0.9870 | 0.9220 | <0.0001 | <0.0001 | 1.0000 | <0.0001 | -- |

**Appendix H4:** Pairwise Comparison Results of **Saccade Offset** Sensitivity Scores

| | | ivt | ivvt | idt | idvt | engbert | nh | remodnav |
|---|---|---|---|---|---|---|---|---|
| ivt | MN | -- | n.s. | n.s. | n.s. | n.s. | n.s. | n.s. |
| | RA | -- | n.s. | ** | ** | n.s. | n.s. | n.s. |
| ivvt | MN | 1.0000 | -- | n.s. | n.s. | n.s. | n.s. | n.s. |
| | RA | 1.0000 | -- | *** | *** | n.s. | n.s. | n.s. |
| idt | MN | 0.1478 | 0.0762 | -- | n.s. | n.s. | n.s. | *** |
| | RA | 0.0014 | 0.0005 | -- | n.s. | n.s. | * | *** |
| idvt | MN | 0.1623 | 0.0850 | 1.0000 | -- | n.s. | n.s. | *** |
| | RA | 0.0011 | 0.0004 | 1.0000 | -- | n.s. | * | *** |
| engbert | MN | 0.8610 | 0.7288 | 0.9017 | 0.9149 | -- | n.s. | * |
| | RA | 0.7340 | 0.5919 | 0.2648 | 0.2381 | -- | n.s. | † |
| nh | MN | 1.0000 | 1.0000 | 0.1688 | 0.1847 | 0.8841 | -- | n.s. |
| | RA | 0.9926 | 0.9711 | 0.0271 | 0.0226 | 0.9850 | -- | n.s. |
| remodnav | MN | 0.5759 | 0.7389 | 0.0001 | 0.0001 | 0.0264 | 0.5378 | -- |
| | RA | 0.8806 | 0.9489 | <0.0001 | <0.0001 | 0.0664 | 0.4346 | -- |

#### Appendix I: HFC-Image Sample-Level Evaluation

We applied the *Sample-Level Evaluation* procedure described above, to the *HFC-image* dataset, using *RA* and *MN* as ground truth annotators. Performance was assessed based on agreement with the GT – measured using *Cohen's Kappa*, *MCC*, and *1-NLD* – and based on fixation sensitivity index ($d'$). Friedman tests revealed significant differences in detector performance for all metrics, regardless of chosen GT annotator.

Pairwise post-hoc comparisons using the Tukey-HSD test showed that the *Engbert* and *NH* detectors significantly outperformed *REMoDNaV*, *I-VT* and *I-VVT*, on all performance metrics. While *I-DT* and *I-DVT* also outperformed these three detectors, the differences were consistently significant only when comparing *I-DVT* with *I-VVT*. In the following pairwise comparison result tables, the top section indicates significance levels, and the bottom section shows the corrected p-values. Significance is denoted as follows:

$$\dagger: p < 0.075, \quad *: p < 0.05, \quad **: p < 0.01, \quad ***: p < 0.001, \quad n.s.: \text{not significant}$$

**Appendix I1:** Sample-Level Evaluation (HFC-Image)

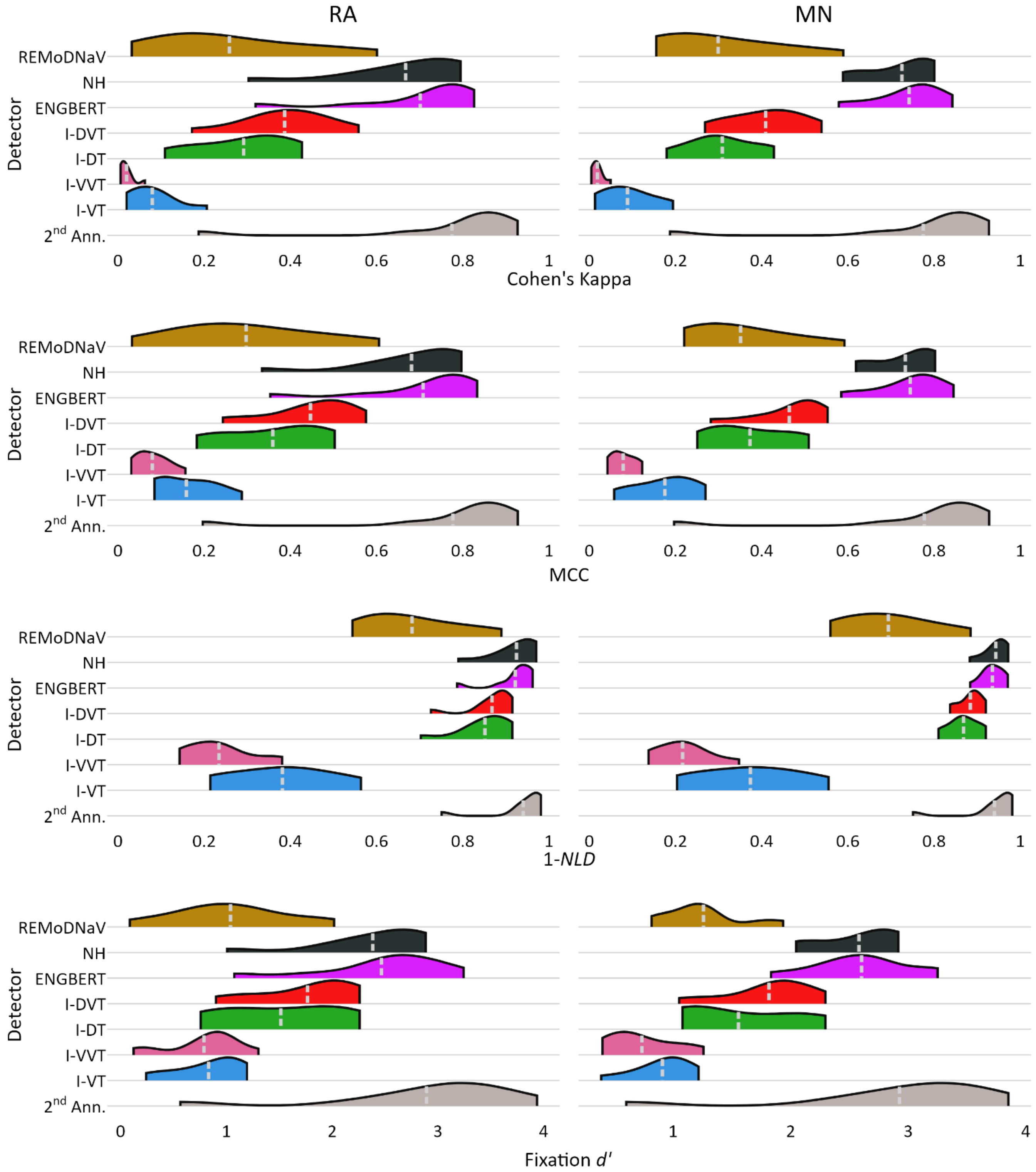


*Note:* Distribution of sample-level performance across recordings of the HFC-Image dataset, with human annotators *RA* and *MN* used as GT (left and right column, respectively), and the 2nd annotator's performance provided for reference (gray violin). The mean of each distribution is denoted by a dashed light-gray line. Across all performance metrics, *NH* and *Engbert's* algorithm perform comparably well or better than the other detectors.

**Appendix I2:** Friedman Test Results

| | **Cohen's Kappa** | | **MCC** | | **1-NLD** | | **Fixation $d'$** | |
|---|---|---|---|---|---|---|---|---|
| | $Q(6)$ | $p$ | $Q(6)$ | $p$ | $Q(6)$ | $p$ | $Q(6)$ | $p$ |
| ***RA*** | $56.0$ | $< 0.001$ | $55.8$ | $< 0.001$ | $58.8$ | $< 0.001$ | $54.2$ | $< 0.001$ |
| ***MN*** | $55.6$ | $< 0.001$ | $55.6$ | $< 0.001$ | $58.8$ | $< 0.001$ | $52.8$ | $< 0.001$ |

**Appendix I3:** Pairwise Comparison Results (*Cohen's Kappa* Scores)

| | | ivt | ivvt | idt | idvt | engbert | nh | remodnav |
|---|---|---|---|---|---|---|---|---|
| ivt | MN | -- | n.s. | n.s. | n.s. | *** | *** | n.s. |
| | RA | -- | n.s. | n.s. | n.s. | *** | ** | n.s. |
| ivvt | MN | 0.9900 | -- | n.s. | * | *** | *** | n.s. |
| | RA | 0.9788 | -- | n.s. | * | *** | *** | n.s. |
| idt | MN | 0.6768 | 0.1969 | -- | n.s. | n.s. | n.s. | n.s. |
| | RA | 0.6528 | 0.1361 | -- | n.s. | n.s. | n.s. | n.s. |
| idvt | MN | 0.2221 | 0.0250 | 0.9933 | -- | n.s. | n.s. | n.s. |
| | RA | 0.2221 | 0.0163 | 0.9949 | -- | n.s. | n.s. | n.s. |
| engbert | MN | 0.0003 | <0.0001 | 0.1700 | 0.5973 | -- | n.s. | n.s. |
| | RA | 0.0008 | <0.0001 | 0.2783 | 0.7232 | -- | n.s. | n.s. |
| nh | MN | 0.0004 | <0.0001 | 0.1929 | 0.6344 | 1.0000 | -- | n.s. |
| | RA | 0.0024 | 0.0000 | 0.4108 | 0.8435 | 1.0000 | -- | n.s. |
| remodnav | MN | 0.7772 | 0.2783 | 1.0000 | 0.9799 | 0.1128 | 0.1300 | -- |
| | RA | 0.8478 | 0.2883 | 0.9999 | 0.9585 | 0.1300 | 0.2178 | -- |

**Appendix I4:** Pairwise Comparison Results (*MCC* Scores)

| | | ivt | ivvt | idt | idvt | engbert | nh | remodnav |
|---|---|---|---|---|---|---|---|---|
| ivt | MN | -- | n.s. | n.s. | n.s. | *** | *** | n.s. |
| | RA | -- | n.s. | n.s. | n.s. | ** | ** | n.s. |
| ivvt | MN | 0.9906 | -- | n.s. | * | *** | *** | n.s. |
| | RA | 0.9887 | -- | n.s. | * | *** | *** | n.s. |
| idt | MN | 0.6886 | 0.2093 | -- | n.s. | n.s. | n.s. | n.s. |
| | RA | 0.7118 | 0.2135 | -- | n.s. | n.s. | n.s. | n.s. |
| idvt | MN | 0.2221 | 0.0258 | 0.9923 | -- | n.s. | n.s. | n.s. |
| | RA | 0.2493 | 0.0283 | 0.9933 | -- | n.s. | n.s. | n.s. |
| engbert | MN | 0.0004 | <0.0001 | 0.1700 | 0.6097 | -- | n.s. | n.s. |
| | RA | 0.0013 | <0.0001 | 0.2733 | 0.7399 | -- | n.s. | n.s. |
| nh | MN | 0.0004 | <0.0001 | 0.1812 | 0.6283 | 1.0000 | -- | n.s. |
| | RA | 0.0027 | <0.0001 | 0.3697 | 0.8303 | 1.0000 | -- | n.s. |
| remodnav | MN | 0.8023 | 0.3091 | 1.0000 | 0.9740 | 0.1049 | 0.1128 | -- |
| | RA | 0.9324 | 0.4967 | 0.9994 | 0.9130 | 0.0880 | 0.1361 | -- |

**Appendix I5:** Pairwise Comparison Results (*1-NLD* Scores)

| | | ivt | ivvt | idt | idvt | engbert | nh | remodnav |
|---|---|---|---|---|---|---|---|---|
| ivt | MN | -- | n.s. | n.s. | n.s. | *** | *** | n.s. |

| | | | | | | | | |
|---|---|---|---|---|---|---|---|---|
| | RA | -- | n.s. | n.s. | n.s. | ** | ** | n.s. |
| ivvt | MN | 0.9945 | -- | * | * | *** | *** | n.s. |
| | RA | 0.9960 | -- | * | * | *** | *** | n.s. |
| idt | MN | 0.2587 | 0.0420 | -- | n.s. | n.s. | n.s. | n.s. |
| | RA | 0.1774 | 0.0266 | -- | n.s. | n.s. | n.s. | n.s. |
| idvt | MN | 0.1183 | 0.0129 | 0.9999 | -- | n.s. | n.s. | n.s. |
| | RA | 0.0903 | 0.0101 | 1.0000 | -- | n.s. | n.s. | n.s. |
| engbert | MN | 0.0008 | <0.0001 | 0.6708 | 0.8601 | -- | n.s. | † |
| | RA | 0.0014 | <0.0001 | 0.8392 | 0.9374 | -- | n.s. | n.s. |
| nh | MN | 0.0003 | <0.0001 | 0.5470 | 0.7668 | 1.0000 | -- | * |
| | RA | 0.0011 | <0.0001 | 0.8071 | 0.9189 | 1.0000 | -- | n.s. |
| remodnav | MN | 0.9298 | 0.5596 | 0.9217 | 0.7721 | 0.0696 | 0.0396 | -- |
| | RA | 0.8937 | 0.5092 | 0.9004 | 0.7772 | 0.1300 | 0.1101 | -- |

**Appendix I6:** Pairwise Comparison Results (*Fixation $d'$ Scores*)

| | | ivt | ivvt | idt | idvt | engbert | nh | remodnav |
|---|---|---|---|---|---|---|---|---|
| ivt | MN | -- | n.s. | n.s. | n.s. | *** | *** | n.s. |
| | RA | -- | n.s. | n.s. | n.s. | ** | ** | n.s. |
| ivvt | MN | 0.9996 | -- | n.s. | n.s. | *** | *** | n.s. |
| | RA | 1.0000 | -- | n.s. | n.s. | ** | ** | n.s. |
| idt | MN | 0.5533 | 0.2783 | -- | n.s. | n.s. | n.s. | n.s. |
| | RA | 0.7003 | 0.5973 | -- | n.s. | n.s. | n.s. | n.s. |
| idvt | MN | 0.2354 | 0.0815 | 0.9990 | -- | n.s. | n.s. | n.s. |
| | RA | 0.3144 | 0.2309 | 0.9979 | -- | n.s. | n.s. | n.s. |
| engbert | MN | 0.0007 | 0.0001 | 0.3471 | 0.6886 | -- | n.s. | † |
| | RA | 0.0041 | 0.0021 | 0.4471 | 0.8212 | -- | n.s. | * |
| nh | MN | 0.0007 | 0.0001 | 0.3415 | 0.6827 | 1.0000 | -- | † |
| | RA | 0.0075 | 0.0040 | 0.5470 | 0.8867 | 1.0000 | -- | * |
| remodnav | MN | 0.9324 | 0.7399 | 0.9937 | 0.9036 | 0.0642 | 0.0624 | -- |
| | RA | 0.9988 | 0.9949 | 0.9421 | 0.6648 | 0.0311 | 0.0499 | -- |

#### Appendix J: Fixation Temporal Alignment (HFC-Image)

Detector's temporal alignment in the *HFC-image* dataset was assessed using the same temporal threshold applied in the main analysis ($|\Delta t| \leq 20\ samples$), equivalent to $66.67ms$. The *Engbert* detector demonstrated the best performance, successfully detecting $99\%$ of fixations with $|RTO| \leq 2.0$ and $RTD \leq 3.5$ samples, surpassing the performance of the 2nd human annotator. The second-best algorithms, *NH* and *REMoDNaV*, detected fixations less accurately, achieving success rates of roughly $80\%$, with $|RTO| \leq 2.5$ and $RTD \leq 4.0$. The worst performing detector, *I-VVT*, had a hit-rate of less than $40\%$ for both fixation onsets and offsets.

**Appendix J1:** Fixation Temporal Alignment Scores

| GT | RA | | | | | | MN | | | | | |
|---|---|---|---|---|---|---|---|---|---|---|---|---|
| | Fixation Onset | | | Fixation Offset | | | Fixation Onset | | | Fixation Offset | | |
| Detector | Hit-Rate | RTO | RTD | Hit-Rate | RTO | RTD | Hit-Rate | RTO | RTD | Hit-Rate | RTO | RTD |
| 2nd Ann. | 93.8% | 0.3 | 3.5 | 93.8% | -0.4 | 2.3 | 94.9% | -0.3 | 3.5 | 94.9% | 0.4 | 2.3 |
| I-VT | 62.5% | 3.5 | 8.7 | 65.1% | -3.1 | 7.5 | 64.5% | 3.3 | 9.0 | 65.2% | -2.6 | 7.2 |
| I-VVT | 38.2% | 2.8 | 10.3 | 38.2% | -4.4 | 8.8 | 39.2% | 3.4 | 10.5 | 39.2% | -3.3 | 9.4 |
| I-DT | 52.9% | -5.7 | 5.0 | 52.2% | 1.7 | 5.7 | 52.8% | -6.1 | 4.5 | 52.8% | 2.5 | 4.3 |
| I-DVT | 55.3% | -5.7 | 4.9 | 55.5% | 2.2 | 5.2 | 55.2% | -6.1 | 4.4 | 56.4% | 2.9 | 4.2 |
| Engbert | **99.0%** | **-1.5** | **3.5** | **98.8%** | **0.9** | **2.5** | **99.0%** | **-2.0** | **2.5** | **99.0%** | **1.3** | **1.7** |
| NH | 77.6% | -2.2 | **4.0** | **76.9%** | **-1.5** | **2.7** | 78.8% | -2.5 | **3.1** | **78.8%** | **-1.1** | **2.5** |
| REMoDNaV | **81.0%** | **-1.7** | 4.3 | 73.3% | **1.5** | 3.9 | **81.1%** | **-2.0** | 3.4 | 71.5% | 2.0 | 3.0 |

#### Appendix K: Fixation Boundary Sensitivity (HFC-Image)

We applied an *Event-Boundary Sensitivity Evaluation* procedure to the *HFC-image* dataset, using the same temporal thresholds as in the main analysis: we calculated fixation onset and offset sensitivity indices ($d'$) for incremental window sizes ($\Delta t \leq 0,1,\dots,20\ samples$), followed by a statistical evaluation for a stringent window size of$|\Delta t| \leq 5\ samples$, equivalent to $16.67ms$. Friedman tests confirmed significant differences in $d'$ scores between detectors for both fixation onsets and offsets, regardless of the GT annotator.

Post-hoc pairwise comparisons using the Tukey-HSD test revealed that the *Engbert* and *NH* detectors performed similarly well for fixation onsets and offsets, significantly outperforming the *I-VT* and *I-VVT* algorithms which showed the poorest sensitivity In the following pairwise comparison result tables, the top section of each table indicates significance levels, while the bottom section displays the corrected p-values. Significance is denoted as follows:

$$\dagger{:}\,p < 0.075, \quad *{:}\,p < 0.05, \quad **{:}\,p < 0.01, \quad ***{:}\,p < 0.001, \quad n.s.{:}\,\text{not significant}$$

**Appendix K1:** Fixation Boundary Sensitivity Index ($d'$) Across Temporal Thresholds Relative to Annotator *RA*

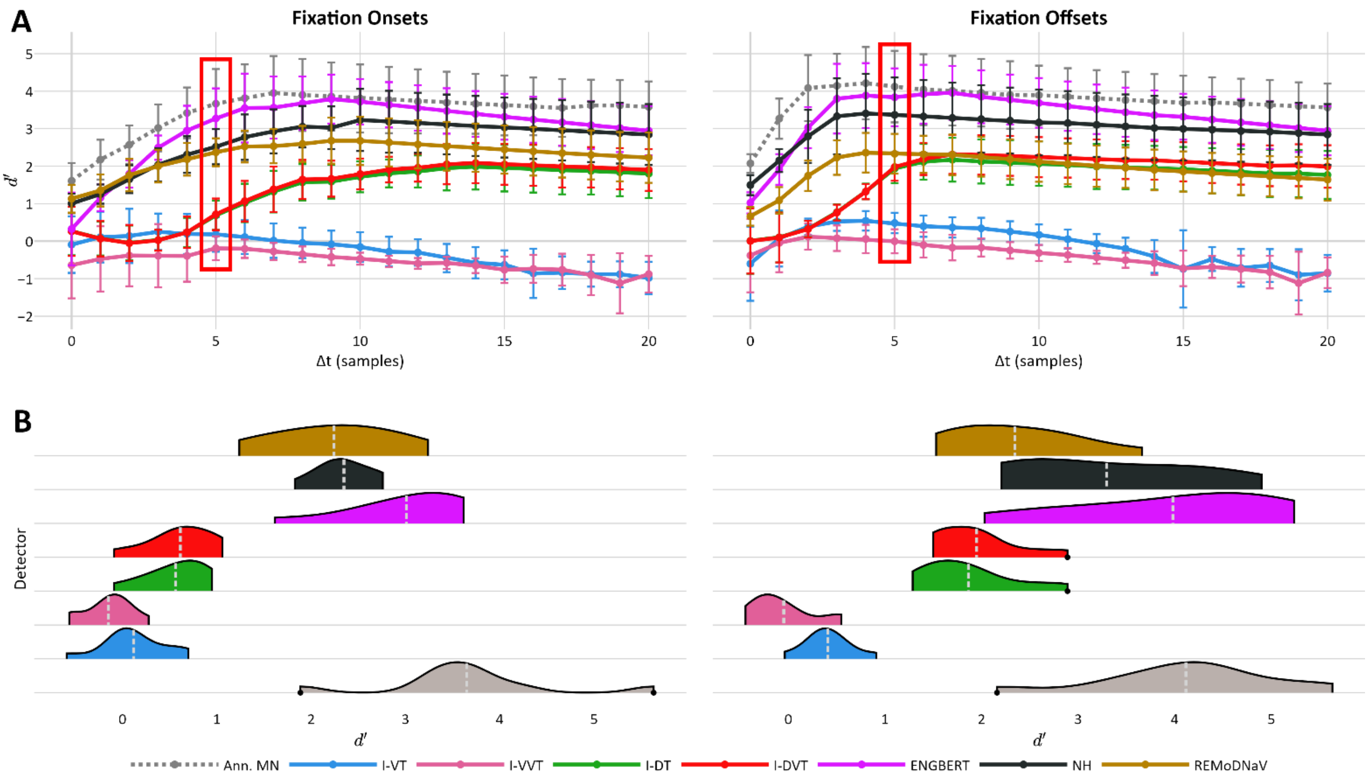


*Note:* Similar to **Figure 4**, this figure shows fixation onset and offset sensitivity scores ($d'$) for each detector, with human annotator *RA* as GT. The top row depicts sensitivity scores across increasing temporal windows ($|\Delta t| = 0,1,\dots,20\ samples$). Each line corresponds to a detector's mean $d'$ across recordings of the *HFC-Image* dataset, and error bars indicate standard deviation. The red rectangle marks the threshold $\Delta t$ used to compare $d'$ scores across detectors, as shown in the bottom row. Sensitivity scores of the second human annotator (*MN*) are also shown for reference (dashed gray line & violin).

**Appendix K2:** Fixation Boundary Sensitivity Index ($d'$) Across Temporal Thresholds Relative to Annotator *MN*

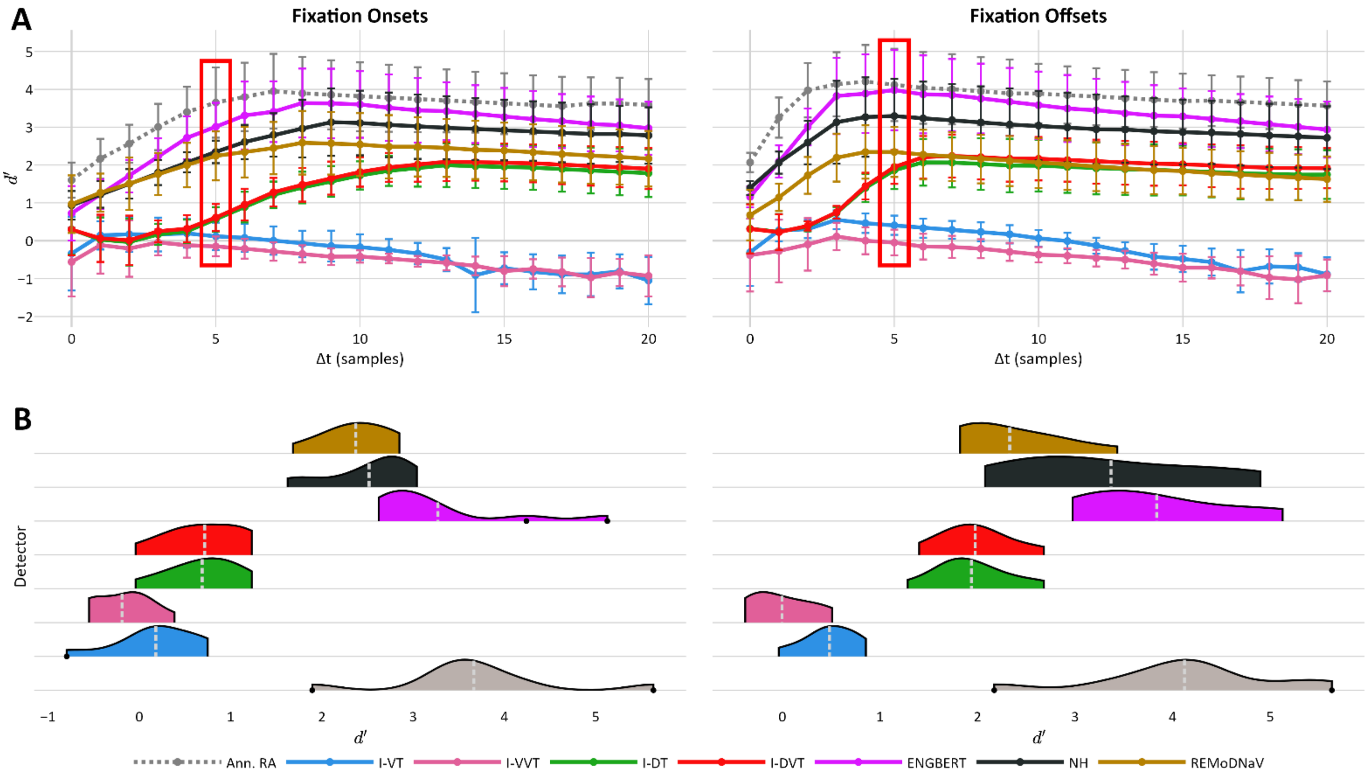


*Note:* Same as **Appendix K1**, but using human annotator *MN* as GT. Sensitivity scores of the second human annotator (*RA*) are also shown for reference (dashed gray line & violin).

**Appendix K3:** Friedman Test Results

| | **Fixation Onset** | | **Fixation Offset** | |
|---|---|---|---|---|
| | $Q(6)$ | $p$ | $Q(6)$ | $p$ |
| ***RA*** | 55.9 | $< 0.001$ | 58.4 | $< 0.001$ |
| ***MN*** | 53.1 | $< 0.001$ | 56.3 | $< 0.001$ |

*Note:* Results of Friedman tests comparing fixation onset and offset sensitivity indices ($d'$) across the seven detectors ($df = 6$), conducted separately using each human annotator (*RA* and *MN*) as ground truth.

**Appendix K4:** Pairwise Comparison Results of **Fixation Onset** Sensitivity Scores

| | | ivt | ivvt | idt | idvt | engbert | nh | remodnav |
|---|---|---|---|---|---|---|---|---|
| ivt | MN | -- | n.s. | n.s. | n.s. | *** | * | * |
| | RA | -- | n.s. | n.s. | n.s. | *** | * | * |
| ivvt | MN | 0.9880 | -- | n.s. | n.s. | *** | *** | ** |
| | RA | 0.9960 | -- | n.s. | n.s. | *** | *** | ** |
| idt | MN | 0.9661 | 0.5941 | -- | n.s. | * | n.s. | n.s. |
| | RA | 0.9603 | 0.6708 | -- | n.s. | * | n.s. | n.s. |
| idvt | MN | 0.9645 | 0.5879 | 1.0000 | -- | * | n.s. | n.s. |
| | RA | 0.9398 | 0.6097 | 1.0000 | -- | * | n.s. | n.s. |
| engbert | MN | 0.0003 | <0.0001 | 0.0238 | 0.0246 | -- | n.s. | n.s. |
| | RA | 0.0003 | <0.0001 | 0.0250 | 0.0341 | -- | n.s. | n.s. |
| nh | MN | 0.0153 | 0.0005 | 0.2563 | 0.2611 | 0.9830 | -- | n.s. |
| | RA | 0.0153 | 0.0009 | 0.2733 | 0.3251 | 0.9810 | -- | n.s. |
| remodnav | MN | 0.0407 | 0.0018 | 0.4258 | 0.4318 | 0.9298 | 1.0000 | -- |
| | RA | 0.0206 | 0.0014 | 0.3197 | 0.3755 | 0.9699 | 1.0000 | -- |

**Appendix K5:** Pairwise Comparison Results of **Fixation Offset** Sensitivity Scores

| | | ivt | ivvt | idt | idvt | engbert | nh | remodnav |
|---|---|---|---|---|---|---|---|---|
| ivt | MN | -- | n.s. | n.s. | n.s. | *** | ** | n.s. |
| | RA | -- | n.s. | n.s. | n.s. | *** | ** | n.s. |
| ivvt | MN | 0.9960 | -- | n.s. | n.s. | *** | *** | * |
| | RA | 0.9975 | -- | n.s. | n.s. | *** | *** | * |
| idt | MN | 0.5973 | 0.1969 | -- | n.s. | n.s. | n.s. | n.s. |
| | RA | 0.5785 | 0.2093 | -- | n.s. | n.s. | n.s. | n.s. |
| idvt | MN | 0.5407 | 0.1628 | 1.0000 | -- | n.s. | n.s. | n.s. |
| | RA | 0.4471 | 0.1330 | 1.0000 | -- | n.s. | n.s. | n.s. |
| engbert | MN | 0.0001 | <0.0001 | 0.1361 | 0.1664 | -- | n.s. | n.s. |
| | RA | 0.0002 | <0.0001 | 0.1851 | 0.2783 | -- | n.s. | n.s. |
| nh | MN | 0.0019 | 0.0001 | 0.4349 | 0.4904 | 0.9981 | -- | n.s. |
| | RA | 0.0024 | 0.0001 | 0.4842 | 0.6159 | 0.9989 | -- | n.s. |
| remodnav | MN | 0.1628 | 0.0234 | 0.9923 | 0.9960 | 0.5470 | 0.8831 | -- |
| | RA | 0.1212 | 0.0186 | 0.9856 | 0.9965 | 0.6945 | 0.9374 | -- |

#### Appendix L: Fixation and Saccade Sensitivity

We computed detection sensitivity indices ($d'$) for fixation and saccade onsets and offsets, for each recording in the *lund2013⁺-image* and *HFC-image* datasets, using both *RA* and *MN* as GT annotators. We applied a temporal window of $\Delta t \leq 5\ samples$ to evaluate $d'$scores.

**Appendix L1:** Detection Sensitivity Index ($d'$) for Fixation Boundaries (*lund2013⁺-image*)

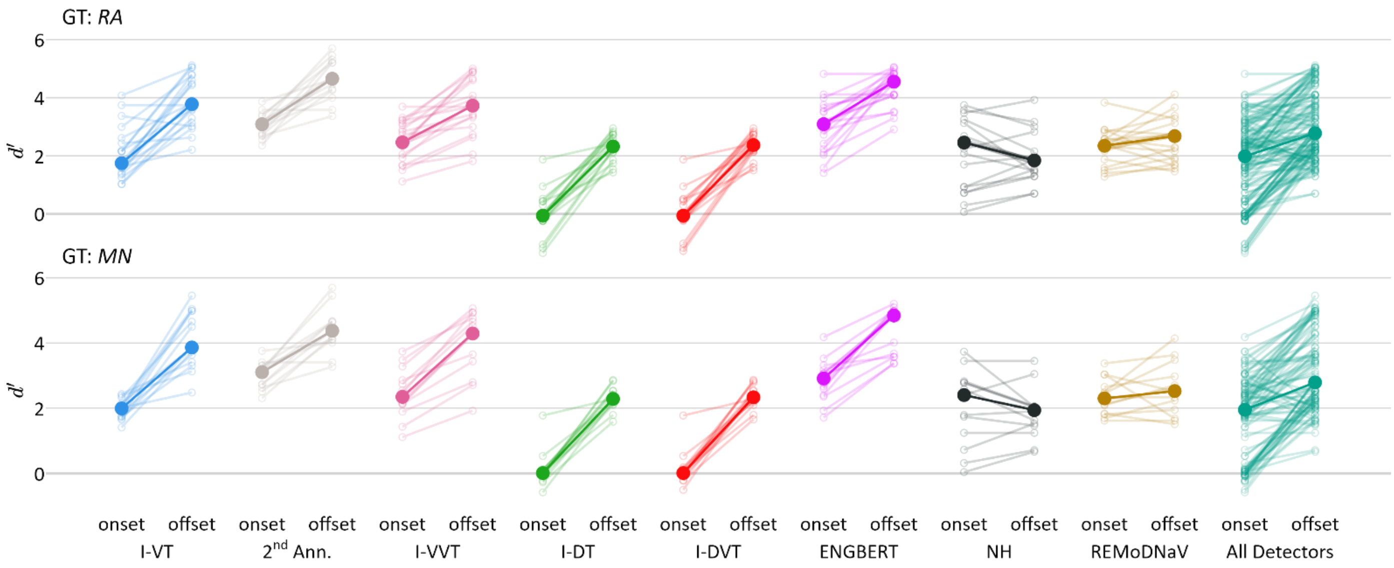


*Note:* Similar to **Figure 6**, this figure shows fixation onset and offset sensitivity indices ($d'$) for each detector, using human annotators *RA* (top row) and *MN* (bottom row) as GT, based on the *lund2013⁺-image* dataset. Sensitivity was computed using a temporal window of $\Delta t \leq 5\ samples$. Small, light-colored circles represent individual recordings, with lines connecting onset and offset $d'$scores from the same recording. Large, opaque circles represent indicate each detector's median $d'$ score. The "All Detectors" column (in green-teal) reflects the overall distribution across all detectors. Results from the second annotator are shown for comparison (in light gray).

**Appendix L2:** Detection Sensitivity Index ($d'$) for Fixation Boundaries (*HFC-image*)

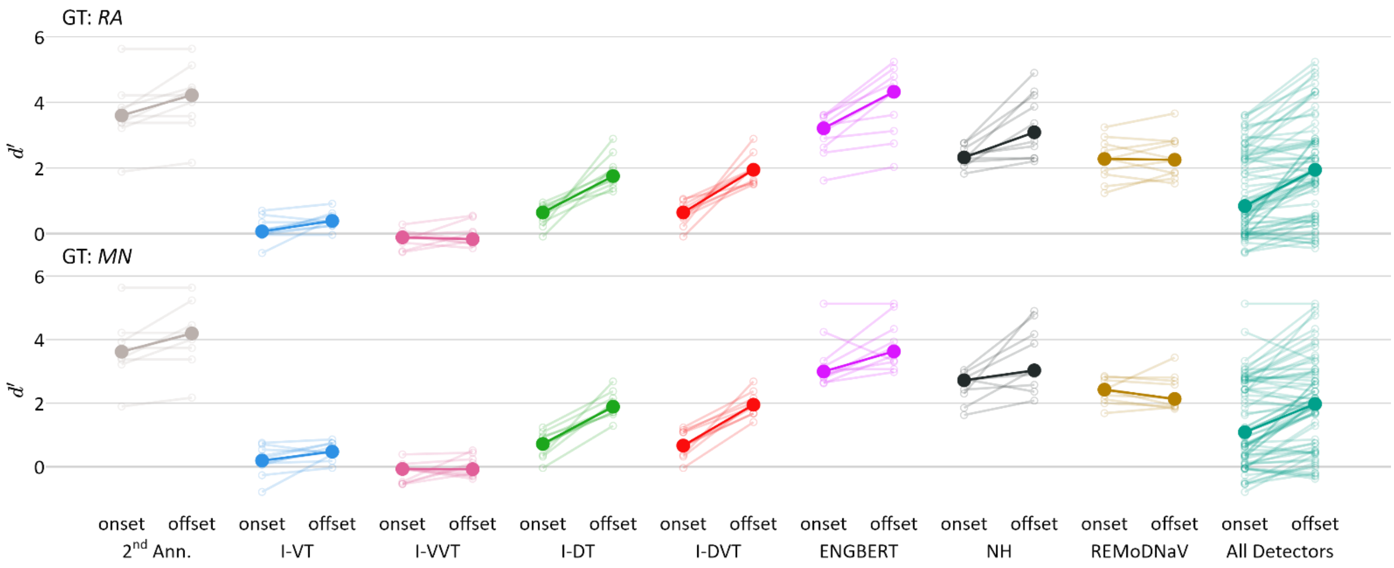


*Note:* Same as **Appendix L1**, but based on the HFC-image dataset.

**Appendix L3:** Wilcoxon Signed-Rank Test Results

We performed a Wilcoxon Signed-Rank test to compare onset and offset $d'$ scores across detectors. Separate tests were conducted for fixations and saccades detected in the *lund2013⁺-image* dataset, and for fixations detected in the *HFC-image* dataset, and performed independently for each GT annotator (*RA* and *MN*).

The following table reports the number of paired samples ($N_{pairs} = N_{recordings} \times N_{detectors}$), the median $d'$ score for onsets and offsets, and the Wilcoxon test statistic ($W$) and $p$-value.

Results indicate a systematic difference in detection sensitivity between onsets and offsets across annotators, event types and datasets.

| | | | **Median $d'$** | | **Wilcoxon Results** | |
|---|---|---|---|---|---|---|
| **Dataset (event type)** | **GT** | $N_{pairs}$ | **onset** | **offset** | $W$ | $p$ |
| ***lund2013⁺-image*** **(saccades)** | **RA** | 140 | 3.2 | 1.9 | 9348 | $< 0.001$ |
| | **MN** | 98 | 3.3 | 1.6 | 4671 | $< 0.001$ |
| ***lund2013⁺-image*** **(fixations)** | **RA** | 140 | 2.0 | 2.0 | 710.5 | $< 0.001$ |
| | **MN** | 98 | 2.8 | 2.8 | 262 | $< 0.001$ |
| ***HFC-image*** **(fixations)** | **RA** | 70 | 0.8 | 1.9 | 161 | $< 0.001$ |
| | **MN** | 70 | 1.1 | 2.0 | 284 | $< 0.001$ |
